# Mechanical morphotype switching as an adaptive response in mycobacteria

**DOI:** 10.1101/2022.04.06.487411

**Authors:** Haig Alexander Eskandarian, Yu-Xiang Chen, Chiara Toniolo, Juan M. Belardinelli, Zuzana Palcekova, Paul Ashby, Georg E. Fantner, Mary Jackson, John D. McKinney, Babak Javid

## Abstract

Invading microbes face a myriad of cidal mechanisms of phagocytes that inflict physical damage to microbial structures. How intracellular bacterial pathogens adapt to these stresses is not fully understood. Here, we report a new virulence mechanism by which mycobacteria alter the mechanical stiffness of their cell surface to become refractory to killing during infection. Long-Term Time-Lapse Atomic Force Microscopy was used to reveal a process of “mechanical morphotype switching” in mycobacteria exposed to host intracellular stress. A “soft” mechanical morphotype switch enhances tolerance to intracellular macrophage stress, including cathelicidin. Genetic manipulation, by deletion of *uvrA*, or pharmacological treatment, with bedaquiline, locked mycobacteria into a “soft” mechanical morphotype state, enhancing survival in macrophages. Our study proposes microbial mechanical adaptation as a new axis for surviving host-mediated stressors.

**One-Sentence Summary:** Bacteria alter their cell surface mechanical properties to increase survival during macrophage infection.

## Main Text

Cell surface elasticity outlines the physical bounds within which microbial life manifests (*1, 2*). Bactericidal stresses drive the biophysical collapse of cell-surface mechanical properties maintaining cell wall integrity, culminating in lysis (*2–4*). Preventing cell lysis is paramount to cell survival, particularly during host pathogenesis when microbes encounter diverse stresses inflicting physical damage. Maintaining a balance of cell mechanical forces is critical to survival. Here, we sought to define the fundamental principles of mycobacterial cell surface mechanical properties to reveal how mycobacteria physically tolerate the stresses of intracellular infection.

Atomic Force Microscopy has previously been used to characterize perturbations of cell wall components at the bacterial cell surface (*2, 3, 5, 6*). More recently, Long-Term Time-Lapse Atomic Force Microscopy (LTTL-AFM) of mycobacteria has defined a synergistic role for cell biomechanics and molecular components influencing fundamental cell processes, such as division site selection, cell cleavage and pole elongation (*3, 4, 7*). We used LTTL-AFM to dynamically characterize the range in cell surface stiffness in *M. smegmatis* (Msm) along the long axis of the cell (Fig. 1A). Growth in Msm consists of pole elongation and subcellular changes in cell surface stiffness (Fig. 1A, B, movie S1, and fig. S1). Elongation results from an expansion of cell volume by addition of nascent cell wall material near the poles (*1, 3, 7*). The elongating mycobacterial cell surface consists of three discrete regions of mechanically distinct material (fig. S1). Most of the cell surface (∼75%), emanating from mid-cell and excluding the poles, is comprised of stably rigid cell surface material (Fig. 1C, figs. S1 and S2A). Sub-polar regions comprise ∼25% of the cell length (∼20% at the old pole and 7% at new pole, pre-New End Take-Off – NETO) and harbor the addition of nascent, non-crosslinked peptidoglycan (*1, 8*) where surface material begins to gradually increase in stiffness (Fig. 1C and figs. S1 and S2A), revealing a process of “mechanical maturation”. Pole elongation dynamics control the shift in mechanically rigid zones towards mid-cell (*7*). These findings expand the description of mycobacterial growth based on pole elongation and mechanical maturation, which is tightly controlled in space and time.

**Fig. 1.**
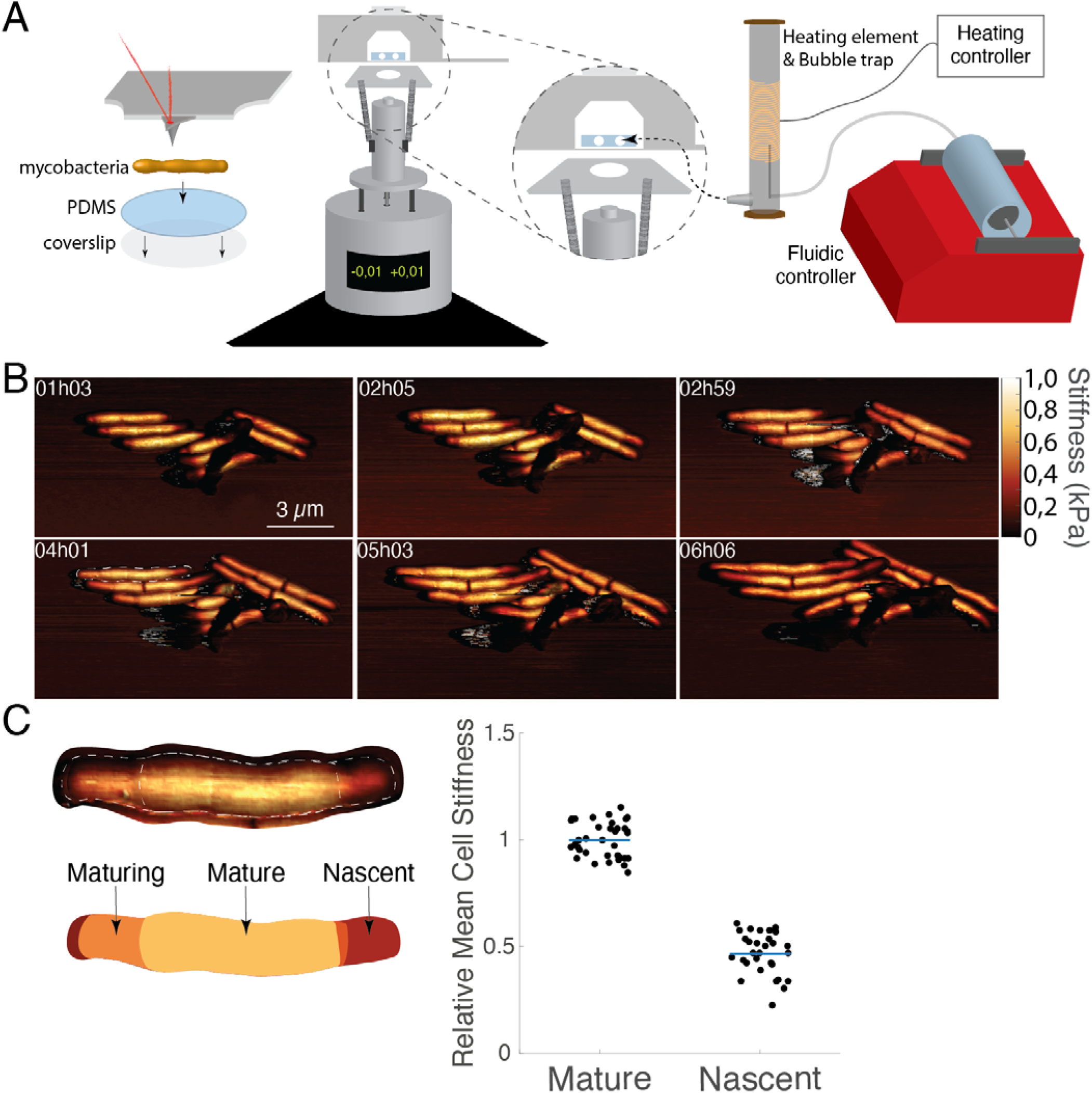
The mycobacterial cell wall controls surface mechanical maturation. **(A)** Schematic representation of the LTTL-AFM imaging system depicting the setup for mycobacterial sample scanning (left) and the integrated heating element and fluidic systems (right) for controlling the environmental conditions while imaging with a Bruker MultiMode 8 scanner. **(B)** Time series of three-dimensional AFM height images overlaid with mechanical stiffness (DMT modulus) images for wild-type *M. smegmatis*. **(C)** The relative cell surface stiffness of nascent cell wall material as compared to mature material. n = 30.

Perturbation of the mycobacterial cell wall, composed of multiple layers that mechanically maintain cell morphology as load-bearing units (*9, 10*), has opposing effects on the cell-surface mechanical state. Cells exhibiting defective peptidoglycan crosslinking due to deletion of L,D– transpeptidases (Δ*ldtAEBCG*+*F*) results in decreased mean cell surface stiffness, culminating in a turgor pressure-driven gradual bulging near the new pole (movie S2) (*1*). By contrast, deleting genes involved in the maturation and export of arabinogalactan and lipoarabinomannan (Δ*Rv1410*-*lprG* and Δ*sucT*) results in increased mean cell surface stiffness, likely by exposing the AFM cantilever to a more load-bearing unit of the cell wall (*11, 12*). Molecular perturbation of biosynthesis processes for discrete layers shifts the mean cell surface stiffness to either “soft” or “hard” mechanical cell states.

Pharmacologic treatment allowed the measurement of dynamic shifts in mean cell-surface stiffness. Mycobacteria treated with isoniazid (INH) gradually drove a two-fold increase in mean cell-surface stiffness and collapsed phenotypic heterogeneity (fig. S3A). Mechanically distinct zones of cell surface material were abolished by INH-induced reductive division and arrest of pole elongation (fig. S2B). Resumed pole elongation resulted in the addition of nascent material undergoing a gradual mechanical maturation returning the mean cell surface stiffness to untreated levels (figs. S3A and S4, movie S3). Treatment with the bacteriostatic cyanide derivative, Carbonyl cyanide m-chlorophenyl hydrazone (CCCP), caused a rapid ∼35% decrease in mean cell surface stiffness (10 kPa min^-1^), which was partially reversed by exchanging the tonicity of the growth medium with water (fig. S3B), suggesting that decreasing mean cell surface stiffness can be mediated by manipulating ATP-dependent control of turgor pressure. Release from CCCP treatment results in the recovery of cell surface stiffness (∼5 kPa min^-1^) prior to the resumption of elongation (figs. S3B, S4, movie S4). The trajectory of mechanical shifts in mean cell surface stiffness are influenced by distinct physical cell properties. Perturbing the inner cell wall results in a decreased mean cell surface stiffness while perturbations to the outer cell wall and cytosolic tonicity drive bacteria into a “hard” mechanical cell state.

*M. smegmatis* represents a technically tractable probe for stress inflicted by macrophages. We used AFM to measure changes to mean cell surface stiffness of Msm bacilli isolated from bone marrow-derived macrophages (Fig. 2, A and B). Following infection, bacilli predominantly had decreased cell-surface stiffness (Fig. 2, B and C). Heterogeneity in cell lengths increased over the course of infection in naïve macrophages among mechanically “soft” bacilli (fig. S5). Some mechanically “soft” bacilli exhibited elongated chains of un-cleaved daughter cells, with two spatially proximal pre-cleavage furrows (Fig. 2B). These chains reflect apparent defects in cleavage reminiscent of peptidoglycan hydrolase mutants or cells exhibiting reduced turgor pressure (*3, 4, 13*). Macrophage stimulation with interferon-gamma (IFNγ) resulted in the emergence of mechanically “hard” bacilli (Fig. 2D). We define these mechanically differentiated, semi-stable cell states within discrete bounds as “mechanical morphotypes”.

**Fig. 2.**
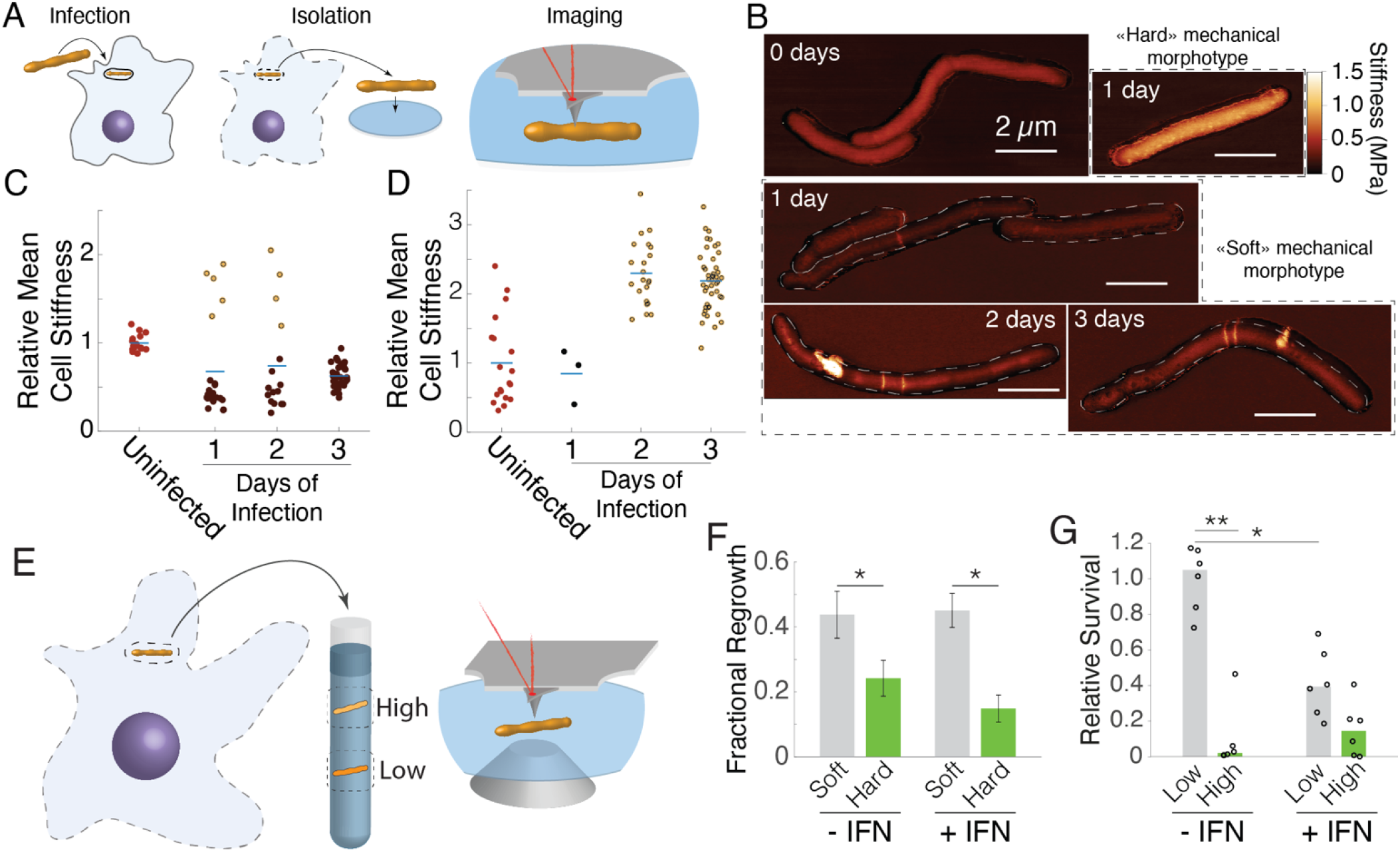
Host macrophage internalization drives selection of distinct mycobacterial mechanical morphotypes. **(A)**, Schematic representation of three critical steps for quantifying surface mechanical properties of macrophage-intracellular mycobacterial cells. **(B and C)** Three-dimensional AFM height images overlaid with stiffness (DMT modulus) images of mycobacterial cells isolated from macrophages during a three-day infection time-course. Normalized relative mean mycobacterial cell stiffness compared to the mean cell stiffness of intracellular mycobacterial cells isolated from macrophages altogether showing the emergence of mechanically “hard” and “soft” cell morphotypes during infection. **(D)** Mycobacteria isolated from macrophages treated with interferon-gamma all exhibited a relative increase in cell surface stiffness. Bar represents mean and dots measurements from individual bacilli. **(E)** Schematic representation of the process by which mechanical morphotypes are isolated from macrophages and enriched by buoyancy fractionation. Enriched mycobacterial morphotypes were analysed for their fractional recovery by quantifying the rate of regrowth using optical fluorescence microscopy **(B)** and by colony forming units (cfu) **(C)**. **(F)** 100 mycobacteria were randomly selected and tracked for 48 hours to quantify whether regrowth would take place. Bars represent the fraction of individuals undergoing regrowth +/- standard error of the mean from three biological replicates (SEM). The “soft” mechanical morphotype exhibited increased regrowth versus the “hard” when isolated from macrophages, independent of cytokine-stimulation. **(G)** Fractional survival as measured by cfu represents values corrected for the number of mycobacteria isolated from macrophages in each sample, showing that cytokine stimulation reduces the yield of sample. \**P* < 0.05, \*\**P* < 0.01 by Student’s T test.

To expand the study of mechanical properties to larger bacterial populations, AFM results were corroborated using isopycnic centrifugation, which enabled fractionation of bacteria based on buoyancy as a surrogate strategy. Samples were fractionated into three buoyancy states, recovering bacilli in “low” (<1.064 g cm^-3^), “middle” (between 1.064 – 1.102 g cm^-3^), and “high” (>1.102 g cm^-3^) buoyancy fractions, based on reported limits in the fractionation of mycobacterial cultures (*14*). Increased cell-surface stiffness corresponded to individuals recovered from a “high” buoyancy fraction and mechanically “soft” bacilli were recovered from a low buoyancy fraction (Fig. S6). Msm isolated from unstimulated macrophages were predominantly recovered from a “low” buoyancy fraction and bacilli isolated from IFNγ-stimulated macrophages were principally recovered from a “high” buoyancy fraction (fig. S7, A-C), recapitulating our findings using AFM. Intracellular *M. smegmatis* isolated from IFNγ-stimulated macrophages treated with bafilomycin A1, an inhibitor of vacuolar maturation, reversed mechanical morphotype enrichment to a “low” buoyancy cell state (figs. S8, S9), supporting a role for phagosomal maturation in the selection of the “hard” mechanical morphotype.

The physiological consequence of mycobacterial differentiation into two distinct mechanical cell states was evaluated by quantifying the rate of resumed growth of individual bacilli representing each discrete mechanical morphotype (Fig. 2E). Intracellular mycobacteria were isolated from lysed macrophages, fractionated by buoyancy centrifugation (Fig. 2E) and the relative recovery rate quantified using LTTL-AFM or plated colony forming units (CFU). Using LTTL-AFM, regrowth of “soft” bacilli was two-fold higher than the “hard” in unstimulated macrophages and three-fold in IFNγ-stimulated macrophages (Fig. 2F). Using colony forming units to quantify relative survival of “low” versus “high” buoyancy fractions, the “low” buoyancy fraction exhibited ten-fold greater survival than the “high” buoyancy fraction in unstimulated macrophages and a two-fold difference in interferon-stimulated macrophages (Fig. 2G).

These results present the possibility that there exists an order of switching from one semi-stable mechanical morphotype to another. We hypothesized that mutants locking *M. smegmatis* into a specific mechanical morphotype state could be isolated with appropriate selection. High density transposon-mutagenized libraries in *M. smegmatis* (*15*) were sorted by buoyancy fractionation to identify genes required for regulating the maintenance of a mechanical morphotype. (fig. S6A). Transposon insertion site frequencies were mapped by sequencing and compared with the unfractionated input library. Conditional essentiality of transposon mutants was identified by the absence of transposon insertions found in “low” (soft) or “high” (hard) buoyancy fractions. These likely corresponded to mutants for which the capacity to switch out of “soft” or “hard” mechanical morphotypes was compromised, resulting in enrichment of the morphotype. We identified a discrete group of genes regulating the “low” buoyancy mechanical morphotype (Fig. 3A). None of the hits identified following stringent post-hoc correction were among highly conserved genes. To extend our findings to other mycobacteria, we asked whether relaxing the post-hoc correction would enable us to identify conserved conditionally essential genes for each discrete mechanical morphotype. We identified many more mutants that appeared to be locked into a “low” buoyancy fraction. By contrast, very few mutants were identified that were locked into a “high” buoyancy fraction (Fig. 3, B, C). The top candidate identified among the “low” buoyancy mechano-morphotype mutants was Δ*Ms3378*, a predicted beta-lactamase (Fig. 3A). Another candidate gene identified in our screen, *uvrA*, is characterized as a member of the *uvrABC* gene system responsible for DNA damage repair and widely conserved in bacteria (*16, 17*) (Fig. 3B). Both Δ*Ms3378* and Δ*uvrA*, exhibited decreased cell surface stiffness by AFM as compared to WT *M. smegmatis* (Fig. 3D, fig. S10, A, B), confirming the validity of our screening approach. Perturbations to genomic stability with mitomycin C resulted in no significant change in the mean cell surface of Msm, despite driving filamentation (*3*) (fig. S11).

**Fig. 3.**
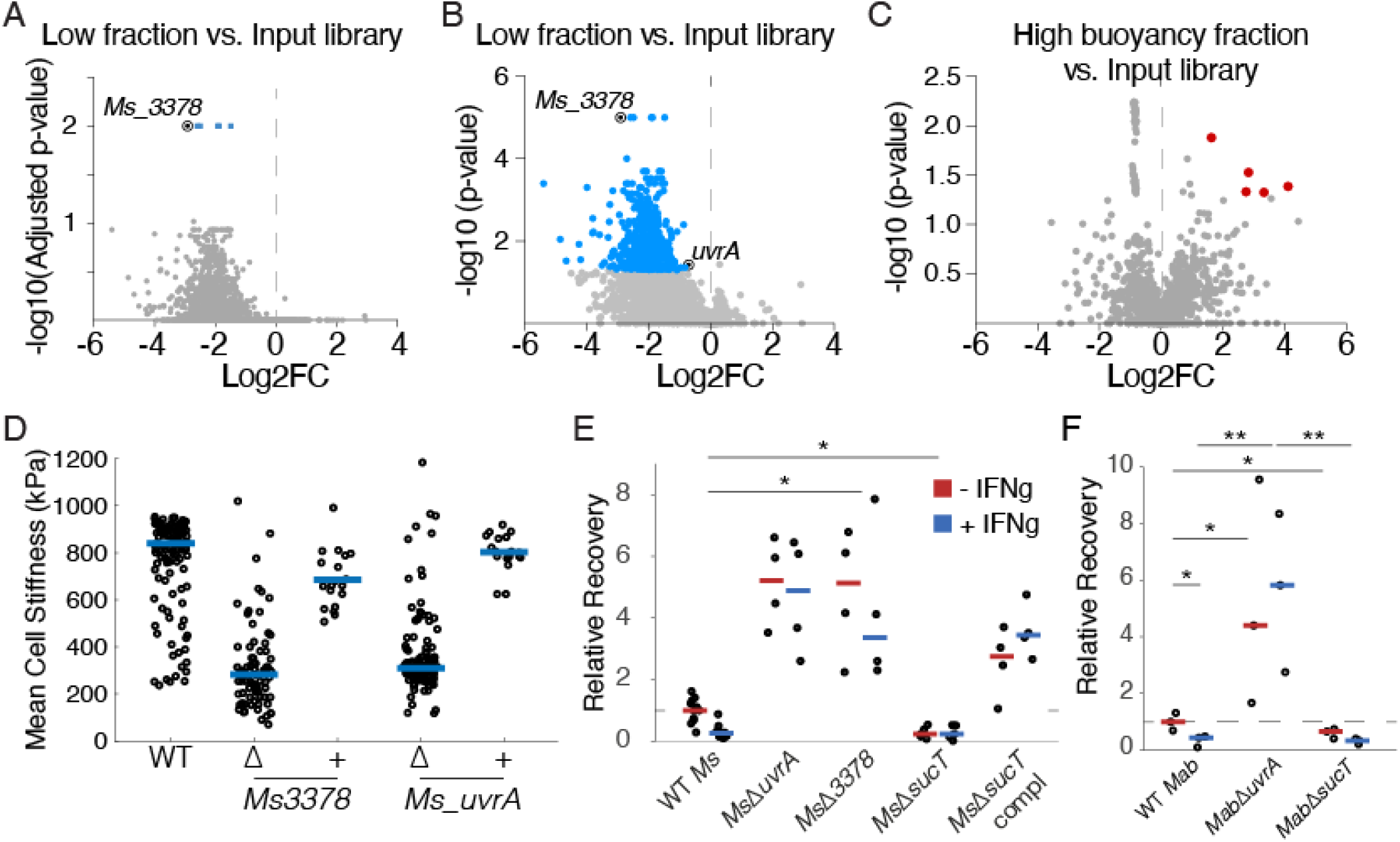
“Soft” mechanical morphotype mutants have enhanced macrophage survival. Volcano plot of genes with different transposon insertions against adjusted p-value (**A**) or p-value (**B**) by re-sampling test for TnSeq analysis of low fraction vs. input library and high fraction vs. input libraries (**C**), genes with differential transposon insertions (see method) are marked in blue and red dots. **(D)** Mean cell surface stiffness, quantified using AFM, of wildtype *M. smegmatis* and “soft” mechano-morphotype mutant strains. **(E and F)** Relative recovery of mechanical morphotype mutants in *M. smegmatis* (**E**) and *M. abscessus* (**F**) following 48-hour infection in macrophages untreated or treated with IFNγ. Bars represent mean. \**P* < 0.05, \*\**P* < 0.01 by Student’s T test.

Buoyancy fractionation enabled us to evaluate the generalizability of mechanical morphotype switching in pathogenic bacteria, which was not possible by AFM due to lack of an instrument in biosafety containment. We interrogated the distribution in mechanical cell states of *M. abscessus* (Mab), a non-tuberculous mycobacterium (NTM) and pathogen. Mab from axenic culture was enriched mostly in the “low” buoyancy fraction (fig. S7D), representing a significant shift as compared with *M. smegmatis*. Smooth and rough colony morphology variants (*18*) did not influence distributions in buoyancy (figs. S7, G, H). Mab recovered from macrophages exhibited the same distributions in buoyancy fractionation as Msm: “low” buoyancy fractions predominated in unstimulated macrophages and “high” buoyancy fractions in IFNγ-stimulated macrophages (fig. S7, E, F), supporting the generalizability of our findings to both non-pathogenic and pathogenic mycobacteria.

We chose *sucT* to evaluate survival of a mutant representing a “hard” mechano-morphotype, a mutant exhibiting elevated mean cell surface stiffness (*12*). We confirmed that Δ*uvrA* and Δ*sucT* had similar phenotypes in Msm and Mab. Buoyancy fractionation of Msm-Δ*sucT* and Mab-Δ*sucT* led to recovery of bacteria predominantly from “high” buoyancy fractions, and this phenotype could be complemented genetically (figs. S12 and S13). As with Msm, Mab-Δ*uvrA* was isolated mostly from “low” buoyancy fractions (fig. S13), confirming that these mutations influence the mechanical state of the cell. In macrophage infection, Mab-Δ*uvrA* was equally enriched in a “low” buoyancy cell state when isolated from both unstimulated and IFNγ-stimulated macrophages. Mab-Δ*sucT* was recovered predominantly from a “high” buoyancy fraction in both unstimulated and IFNγ-stimulated macrophages (fig. S14).

Do mechanical morphotype mutants show altered survival during macrophage infection? The “soft” mechano-morphotype mutants, Msm-Δ*uvrA* and Msm-Δ*Ms3378*, both had increased survival in macrophages, particularly in IFNγ-stimulated macrophages with a ∼6-fold and >20-fold increase, respectively, as compared to wildtype (Fig. 3E). Survival of Msm-Δ*sucT* was attenuated compared with wildtype in both unstimulated and IFNγ-stimulated macrophages (Fig. 3E). Similarly, Mab-Δ*uvrA* isolated from unstimulated macrophages showed 20% increased survival as compared to wildtype and a >5-fold increase upon IFNγ-stimulation (Fig. 3F). Mab-Δ*sucT* had two-fold decreased survival during infection of unstimulated macrophages as compared to wildtype, a host cell condition selecting for the predominance of a “soft” mechanical cell state (Fig. 3F). In IFNγ-stimulated macrophages, which drives selection of a “hard” mechanical cell state (fig. S8F), wildtype *Mab* and Mab-Δ*sucT* had similar rates of recovery (Fig. 3F).

Does altering the mechanical state of wildtype Mab influence survival within macrophages? Macrophages were infected with Mab pre-fractionated into “high” and “low” buoyancy fractions. “Low” buoyancy fractions of Mab showed ∼5-fold increased survival in unstimulated macrophages and >10-fold increased survival in IFNγ-stimulated macrophages, as compared with pre-fractionated “high” buoyancy morphotypes (Fig. 4A). Sub-inhibitory concentrations of antibiotics driving mechanical morphotype switching was used as an orthogonal method to pre-condition wildtype Mab. INH was used to produce “hard” bacilli and bedaquiline (BDQ), an ATPase inhibitor (like CCCP) and clinically relevant antibiotic for the treatment of *M. tuberculosis*, was used to generate “soft” mechanical morphotype bacilli. BDQ-conditioned bacilli exhibited ∼2-fold increased survival in unstimulated macrophages and greater than 5-fold increased survival in IFNγ-stimulated macrophages as compared to both untreated and INH-conditioned bacilli (Fig. 4, B and C). Our results suggest that the impact of antibiotics on mechanical morphotype switching, and hence macrophage survival, may be an important consideration in treating mycobacterial infections.

**Fig. 4.**
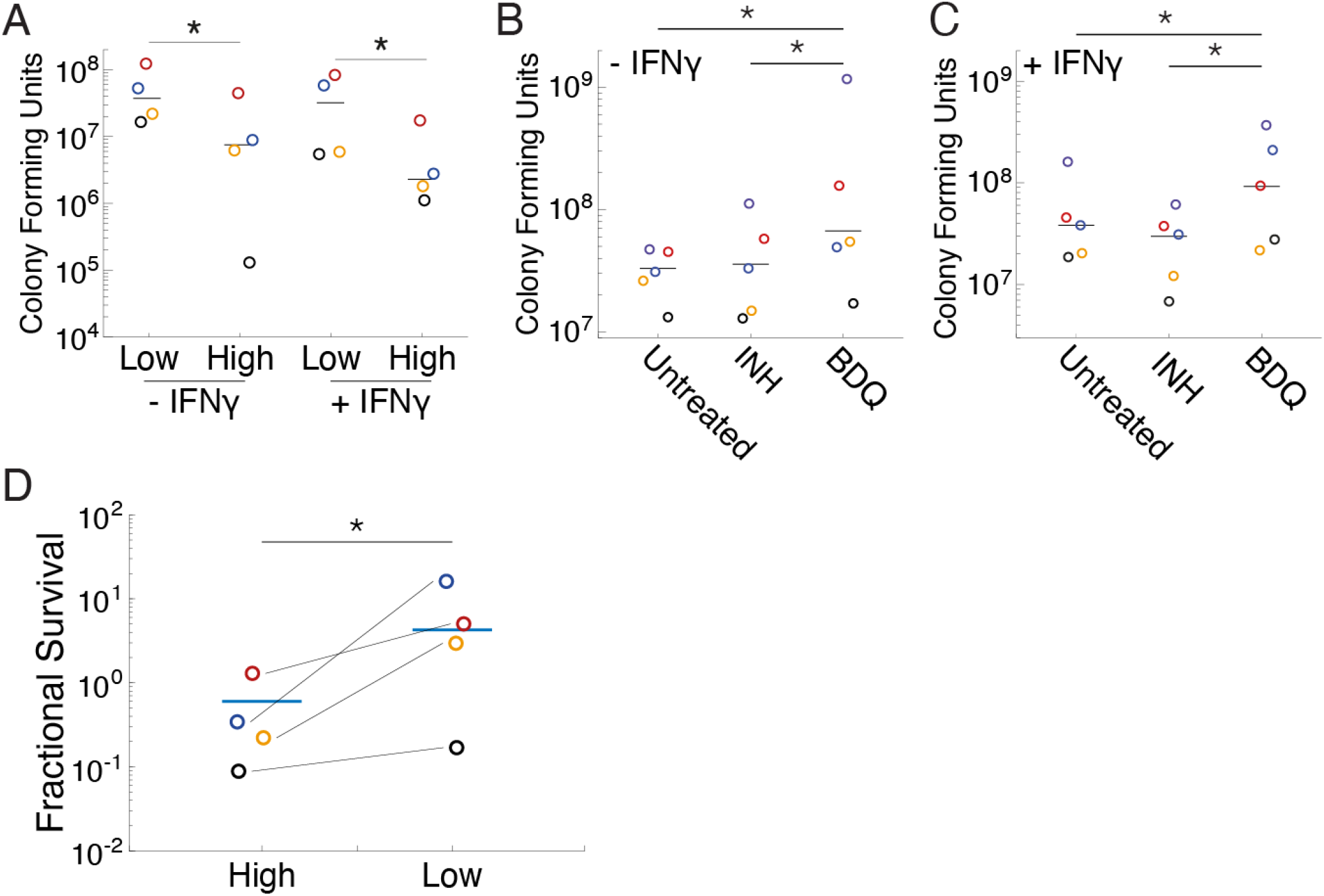
Phenotypic conditioning of mechanical state alters survival of *M. abscessus* to macrophage-derived stress. Mab was preconditioned by buoyancy fractionation (**A**) or sub-MIC antibiotic treatment (**B**) and (**C**) into “high”-buoyancy (hard) and “low”-buoyancy (soft) mechanical morphotypes and then used to infect macrophages (MOI = 10 bacteria per macrophage) for 48 hours. Survival was quantified by harvesting bacteria from macrophages and quantifying colony forming units. *p< 0.05 by Student’s t-test. (**D**) Mechanical morphotypes in Mab, isolated by buoyancy fractionation, were subjected to the antimicrobial peptide, cathelicidin (LL37). Fractional survival of LL37-treated (5µg/ml) versus untreated pre-fractionated Mab mechanical morphotypes. Wilcoxon signed rank pair test p-value = 0.0246. Each colored dot represents a separate experiment and bars represent the mean of the experimental replicates.

The antimicrobial peptide, cathelicidin (LL37), contributes to macrophage-mediated killing of intracellular pathogens, including mycobacteria (*19*) through physical damage. Do mechanical morphotype mutants exhibit differences in survival to LL37-treatment? Wildtype Mab was pre-fractionated by buoyancy centrifugation into “low” and “high” buoyancy fractions and subsequently treated with LL37. The “low” buoyancy fraction exhibited ∼7-fold increased tolerance to LL37-treatment as compared to the “high” buoyancy fraction (Fig. 4D). These results suggest that the “low” buoyancy fraction (“soft” mechanical morphotype) represents an innate immune stress-tolerant phenotypic state in mycobacteria.

Previous studies outline limits in the mechanical variation of bacterial cell wall integrity and cytosolic tonicity (*2, 14, 20-22*), both of which are fundamental cell properties that can influence the impact of bactericidal stress. We demonstrate that mycobacterial cell-surface mechanical stiffness is modulated to optimize survival to cidal stress and can be influenced by genes widely conserved in prokaryotes. Mechanical adaptation might represent a fundamental cell process in all bacteria harnessed to respond to changing environmental conditions. Divergent trajectories of mechanical switching to “hard” and “soft” cell states raises the possibility for bacilli to respond to diverse stresses. Our findings fundamentally expand the understanding of mycobacterial virulence to include mechanical adaptation as an important driver of survival during macrophage infection.

## Acknowledgements

We thank Jeff Cox and Hesper Rego for critical reading of the manuscript.

## Author Contributions

HAE, YC, CT, and BJ conceived of experiments. CT and HAE conducted macrophage infections and optical fluorescence microscopy. YC and HAE conducted transposon mutagenesis and YC analysed TnSeq data. JB produced and provided knockout mutants in *Msm* and *Mab*. PA provided access to AFM system at LBNL. HAE conducted AFM imaging, image processing, and data analyses. JB, ZP, and MJ provided genetic mutants in *Msm* and *Mab*. JDM provided funds and infrastructure at EPFL. BJ provided funds and infrastructure at UCSF. HAE and BJ organised, wrote, and edited manuscript. HAE, YC, CT, BJ, ZP, PA, MJ, JDM and BJ all provided critical edits to the manuscript.

## Data availability

Tnseq raw datasets are available at: https://figshare.com/s/221d724865faa6deb7e4. Select LTTL-AFM datasets are available at: https://figshare.com/s/4a63d8e2186f8246077d. Raw experimental data supporting the findings of these studies are available from the corresponding author upon request.

## Supplementary Materials

### Materials and Methods

#### Bacteria

*Mycobacterium smegmatis* mc^2^155 (wild-type) and derivative strains and *Mycobacterium abscessus* (ATCC 19977) and derivative strains were grown in Middlebrook 7H9 liquid medium (Difco) supplemented with 0.5% albumin, 0.2% glucose, 0.085% NaCl, 0.5% glycerol, and 0.5% Tween-80. Cultures were grown at 37°C to mid-exponential phase (optical density at 600nm (OD_600_) of ∼0.5). Aliquots were stored in 15% glycerol at −80°C and thawed at room temperature before use. The Δ*uvrA* strains were made by allelic exchange with an unmarked in-frame deletion of the *uvrA* gene in both *M. smegmatis* and *M. abscessus*. The *attB*-integrating plasmid expressing a *uvrA*-*wasabi* fusion was used to complement the gene *uvrA* into the *Ms*Δ*uvrA* strain. Construction of the *M. smegmatis sucT* complementation strain was described previously (*12*). Chemico-mechanical manipulation of mycobacteria was conducted by treatments with isoniazid (INH) (Sigma) at 10 µg ml^-1^ (2x minimum inhibitory concentration (MIC)) or 5 µM carbonyl cyanide-m-chlorophenylhydrazone (CCCP). Cathelicidin (LL37) (ThermoFisher) at 0.5 µg ml-1.

#### M. abscessus ATCC 19977 uvrA KO

Deletion of *uvrA* (*MAB_2315*) from *M. abscessus* ATCC 19977 was carried out using the ORBIT system described by Murphy, *et al.* (*23*). Briefly, *M. abscessus* was transformed with plasmid pKM444 expressing the Che9c phage RecT annealase and the Bxb1 phage integrase under control of the inducible pTet promoter. A 20ml culture of *Mabs* pKM444 as grown at 37C until OD=0.5, anhydrotetracycline was added at a final concentration of 500ng/ml to induce expression and cells were further incubated for 4h until OD=1. At this point, cells were harvested, washed 3 times with 10% glycerol and resuspended in 2ml of 10% glycerol. An aliquot of 380ml of cells was co-transformed with 200ng of payload plasmid pKM496 (Zeo^R^) plus 1mg of ORBIT oligo and incubated O/N with shaking at 37°C to let them recover. Cells were collected and spread on 7H11 ADC + Zeo (100mg/ml); plates were incubated for 5 days at 37°C and colonies were picked and checked by PCR to verify the recombinants.

#### Transposon library construction, genomic DNA Extraction, and sequencing

Transposon library after buoyancy centrifugation was collected and resuspended in 400ul 10mM Tris (pH=9). After beads-beating, genomic DNA was extracted by Phenol-Chloroform method. DNA concentration was measured and quantified by Nanodrop and Qubit. For building transposon sequencing library, approximately 5ug genomic DNA was resuspended in 150ul TE buffer and transferred to a Covaris tube, Genomic DNA was disrupted to 200-500bp size range by sonication with the following parameters: duty cycle (10%), intensity (4), cycles/burst (200), time (80s). The fragmented genomic DNA was size-selected and purified by AMPure XP beads. The fragmented genomic DNA was further subjected to end repair and dA tailing. Annealed adapter was ligated to the dA tailed fragmented genomic DNA and the linked ligated DNA fragment was used as template of 1^st^ round nested PCR to amplify fragments containing adaptor and transposon junction. Indexed barcoded sequencing and illumina sequencing adaptor was added by 2^nd^ nested PCR. All sequence libraries were examined by Agilent 2100 Bioanalyzer and subjected to next generation sequencing.

#### Transposon mutagenesis

*Mycobacterium smegmatis* mc^2^155 strain was grown to stationary phase (OD>6) in 50ml of 7H9 growth medium. Bacterial cultures were washed and resuspended in 5ml MP buffer (50mM Tris, 150mM NaCl, 10mM MgSO4, 2mM CaCl2). To transduce bacteria with MycoMarT7 phage, approximately 10^11^ plaque forming units of phage (PFU) was added to the bacterial suspension in MP buffer and incubated at 37°C for 4h. Immediately after transduction, ∼300-400ul of the transduction mixture was plated on 15-cm LB agar plates, containing 20ug/ml kanamycin and 0.1% Tween80. After 3 days, library size was determined, and bacteria was scrapped and stored in 7h9 medium plus 15% glycerol as library stock. The transposon library was made in triplicate. The transposon library was cultured to an OD_600nm_ of 0.8 and 1ml of sample was loaded onto 10ml of stock isotonic percoll medium, with buoyant density beads as fiducial markers. Buoyancy centrifugation was conducted at 18°C, and spinning at 20k rpm (∼50,000 g), for 1h20m. Three buoyancy fractions were isolated: “high” (>1.02 g cm^-3^, <1.064 g cm^-3^), “middle” (>1.064 g cm^-3^, <1.102 g cm^-3^), and “low” (>1.102 g cm^-3^). Three biological replicates of the buoyancy centrifugation were conducted for the transposon library made in triplicate each of the three transposon libraries: 3 (libraries) x 3 (buoyancy centrifugation experiments) x 3 (buoyancy fractions) = 27 individual samples.

#### Transposon mapping and analysis

Reads processing and TA loci mapping were performed through software TRANSIT (*24*). Loci that were differentially disrupted by transposon were analysed using resampling test in TRANSIT with default parameter. Different buoyancy fractions were compared with input libraries and genes that are over-represented <u>(log2FC < −1,</u> <u>adjusted p-value < 0.05)</u> and under-represented <u>(Log2FC >1, p-value < 0.05)</u> were plotted.

#### ORBIT oligo for uvrA KO

CGGTTCACCAACGGCGGCGTCAGTCATGACTGCCACCCTAGACCGGAGTGACAACCTTCCTGGTCCGCGCGGTTTGTACCGTACACCACTGAGACCGCGGTGGTTGACCAGACAAACCCGCGCCCCGCACGATCAGACGGTCGGCCACCGGTCTCCTTTCACACTGCCCTATGCAGGTGTTTTCGCGT

#### Cell culture and Infection

Bone marrow derived macrophages (BMDMs) were differentiated from cryopreserved bone marrow stocks extracted from femurs of 8-week-old C57BL/6 mice. After cultivation for 7 days in Petri dishes in BMDM differentiation medium (DMEM with 10% FBS, 1% sodium-pyruvate, 1% GlutaMax and 20% L929-cell-conditioned medium as a source of granulocyte/macrophage colony stimulating factor) adherent cells were gently lifted from the plate using a cell scraper, resuspended in BMDM culture medium (DMEM with 5% FBS, 1% sodium-pyruvate, 1% GlutaMax and 5% L929-cell-conditioned medium) and seeded on the plate used for the the experiment. For infection, 1 ml of *M. smegmatis* or *M. abscessus* culture at OD_600_ 0.4-0.8 was pelleted, resuspended in 200 μl of BMDM culture medium and passed through a 5 μm filter to eliminate bacterial aggregates. The resulting single-cell suspension was used to infect BMDMs at an MOI of 1:1. After 4 hours of infection, macrophages were washed extensively to remove extracellular bacteria and incubated with fresh macrophage medium. All the incubations were performed at 37°C, 5% CO_2_. When required, 100 U/ml IFNγ was added to the macrophage medium 16 hours before infection and kept during the experiment. At selected time-points post infection macrophages were lysed with 0.5% Triton X-100 in PBS for 5 minutes to collect the intracellular bacteria. The cell lysate was pelleted, washed twice with PBS and the pellet was resuspended in 7H9 for further microscope imaging.

RAW264.7 macrophages were infected with *M. abscessus* or *M. smegmatis* (MOI 10:1) for 30 minutes before washing cells and adding gentamicin (50 µg/ml) for 1 hour to remove extracellular bacilli. When required, IFNγ (100 U/ml) was added to macrophages 12 hours prior to infection and kept during the experiment. At 48 hours of infection, macrophages were thoroughly washed with fresh medium and lysed with 0.5% Triton X-100 in PBS for 5 minutes to collect intracellular bacilli. The cell lysate was pelleted, washed twice with PBS to remove detergent, and samples either serial diluted to plate for CFU or loaded onto standard isotonic Percoll to fractionate differentially buoyant bacilli by buoyancy gradient centrifugation.

#### Optical Fluorescence Microscopy

Bacteria extracted from macrophages were seeded on a 35 mm Ibidi μ-dish and imaged for up to 24 hours at 1-hour intervals to check for growth recovery. Infected BMDMs were imaged for up to 72 hours at 1-hour intervals. All the microscopy images were acquired on a DeltaVision microscope equipped with an enclosure maintaining the temperature of the sample at 37°C, FITC (Ex 490/20, Em 525/36) and TRITC (Ex 555/25, Em 605/52) filters and 60x or 100x objectives. Host cells and bacteria were respectively identified in bright-field and fluorescence images. Multiple XY fields were acquired in parallel. For microscope experiments with macrophages, samples were maintained in a humidified stage-top incubator connected to a gas mixer (Okolab) supplying air mixed to 5% CO_2_. For some experiments, macrophages imaged by time-lapse microscopy were lysed by replacing the macrophage medium with 0.5% Triton X-100 in PBS using custom-made tubing connected to the lid of the macrophage dish. After 5 minutes, cells were washed gently with PBS and then incubated with 7H9 medium supplemented with 100 ng ml^-1^ calcein-AM to identify live, enzymatically active cells. Bacteria sticking to the debris of the lysed cells were imaged by time-lapse microscopy to track their regrowth.

#### Image analysis

The microscopy images and time-series were analysed using the FIJI software from the ImageJ package (*25*). Macrophage infection status and viability were monitored by visual analysis of the time-lapse bright-field image series. Regions of interest corresponding to individual macrophages were manually drawn onto the bright-field images and transferred to fluorescence images. A manual threshold was set on the fluorescent channel to segment the bacteria. The area above the threshold for each region of interest was measured for each time-point and used as a proxy for the number of intracellular bacteria per cell. The bacterial growth rate of each individual intracellular micro-colony was calculated fitting an exponential curve was fitted to the measured fluorescent areas. For individual bacteria extracted from lysed macrophages, regrowth and staining were manually analysed through visual inspection of the microscope image series.

#### AFM imaging

Coverslips were prepared as previously described (*3*). Polydimethylsiloxane (PDMS) (Sylgard 184, Dow Corning) at a ratio of 15:1 (elastomer:curing agent) and cut 1:10 with Hexane to reduce 10-fold the spin-coated layer, while equally increasing the hydrophobicity of the surface. Aliquots of mycobacteria isolated from axenic culture or from infection of macrophages were filtered through a 0.5 µm pore size PVDF filter (Millipore) to remove cell clumps and enrich single cells. Aliquots were deposited on the hydrophobic surface of a PDMS-coated coverslip. 7H9 growth medium was supplied. Where indicated, antibiotic was added to the growth medium. The medium was maintained at 37°C using a custom-made heating element within the sample space and a TC2-80-150 temperature controller (Bioscience tools). Bacteria were imaged by peak force tapping using a Nanoscope 5 controller (Veeco Metrology) at a scan rate of 0.25 – 0.5 Hz and a maximum Z-range of 5 µm. A ScanAsyst fluid cantilever (Bruker) was used. Continuous scanning provided snapshots at 2-30 min intervals. Height, peak force error, adhesion, dissipation, deformation modulus and log modulus were recorded for all scanned images. Peak force error yields a fine representation of the height on the order of 10 nm in the Z-axis; this is computed as the difference between the peak force setpoint and the actual value. Images were processed using Gwyddion (Department of Nanometrology, Czech Metrology Institute). ImageJ was used for extracting bacterial cell surface height and modulus values and generating dynamic and quantitative mean values of the mechanical properties of individual cells.

#### Correlated fluorescence and AFM

Correlated fluorescence and AFM images were acquired as described previously (*3, 26*). Briefly, fluorescence images were acquired with an electron-multiplying charge coupled device (EMCCD) iXon Ultra 897 camera (Andor) mounted on an IX71 inverted optical microscope (Olympus) equipped with an UAPON100XOTIRF x100 oil immersion objective (Olympus) with the x2 magnifier in place. Illumination was provided by a monolithic laser combiner (MLC) (Agilent) using the 488 or 561 nm laser output coupled to an optical fibre with appropriate filter sets: F36-526 for Calcein-AM and F71-866 for mCherry-Wag31 or cytosolic RFP. The AFM was mounted on top of the inverted microscope and images were acquired with a customised Icon scan head (Bruker) using ScanAsyst fluid cantilevers (Bruker) with a nominal spring constant of 0.7 N m^-1^ in peak force tapping mode at a setpoint <2 nN and typical scan rates of 0.5 Hz. The samples were maintained at 37°C in 7H9 growth medium heated by a custom-made coverslip heating holder controlled by a TC2-80-150 temperature controller (Bioscience tools).

#### Cell measurements

Fundamental physiological cell measurements of both cell dimensions and mechanical surface rigidity were made as described previously (*1, 3, 4, 11, 12*).

##### Cell growth measurements

Cell growth herein describes a process of cell expansion, which is defined as a function of both cell elongation along the long axis of the cell and surface mechanical maturation (change in cell surface stiffness) upon the addition of nascent cell wall material. Cell length was measured as per (*3, 4, 7*). Mechanically distinct cell growth zones were segregated into three spatially discrete zones based on the dynamic and mechanical state of the surface: 1) the nascent growth zone consists of mechanically “soft” material typically localised nearest the poles in mycobacteria, 2) a maturation zone consists of the space adjacent to the nascent zone where surface material is dynamically increasing in mechanical rigidity, and 3) a mechanically mature zone where surface material mechanical properties has ceased to increase.

##### Mean cell surface stiffness

Mycobacterial cell surface stiffness is averaged over the relatively flat surface along the longitudinal midline of the cell surface, which is probed by AFM imaging in peak force tapping mode and interpreted as the Young’s modulus as per the Derjaguin-Muller-Toporov (DMT) model. The absolute values can vary from one experiment to the next, as a function of variations in the biophysical properties of the fluid medium (temperature and fluid density). Therefore, the mean cell surface rigidity is interpreted as a relative measure comparing the bacterium to the modulus of the PDMS-coated coverslip sample surface (fig. S14).

##### Buoyant density

Buoyancy fractionation was conducted by centrifuging mycobacteria at high speeds in a viscous, gradient-forming medium. Stock isotonic percoll medium (SIP) is prepared by mixing 9ml Percoll with 1ml 0.15 M NaCl. 10 ml of SIP was loaded into thin wall polypropylene centrifuge tubes (14 x 89 mm) (Beckman Coulter) along with 1 ml of bacterial sample and 50 µl of buoyancy centrifuge beads. Mycobacteria are centrifuged at 20,000 rpm (∼52,000 g) for 1 hour and 20 minutes using a Beckman Coulter Optima L-90K Ultracentrifuge and a SW40Ti swinging bucket rotor. Buoyant density beads (Cospheric) at 1.02 g cm^-3^, 1.064 g cm^-3^, 1.08 g cm^-3^, and 1.102 g cm^-3^ were used to distinguish “high”, “middle” and “low”-buoyancy fractions of mycobacteria.

---

**Fig. S1.**
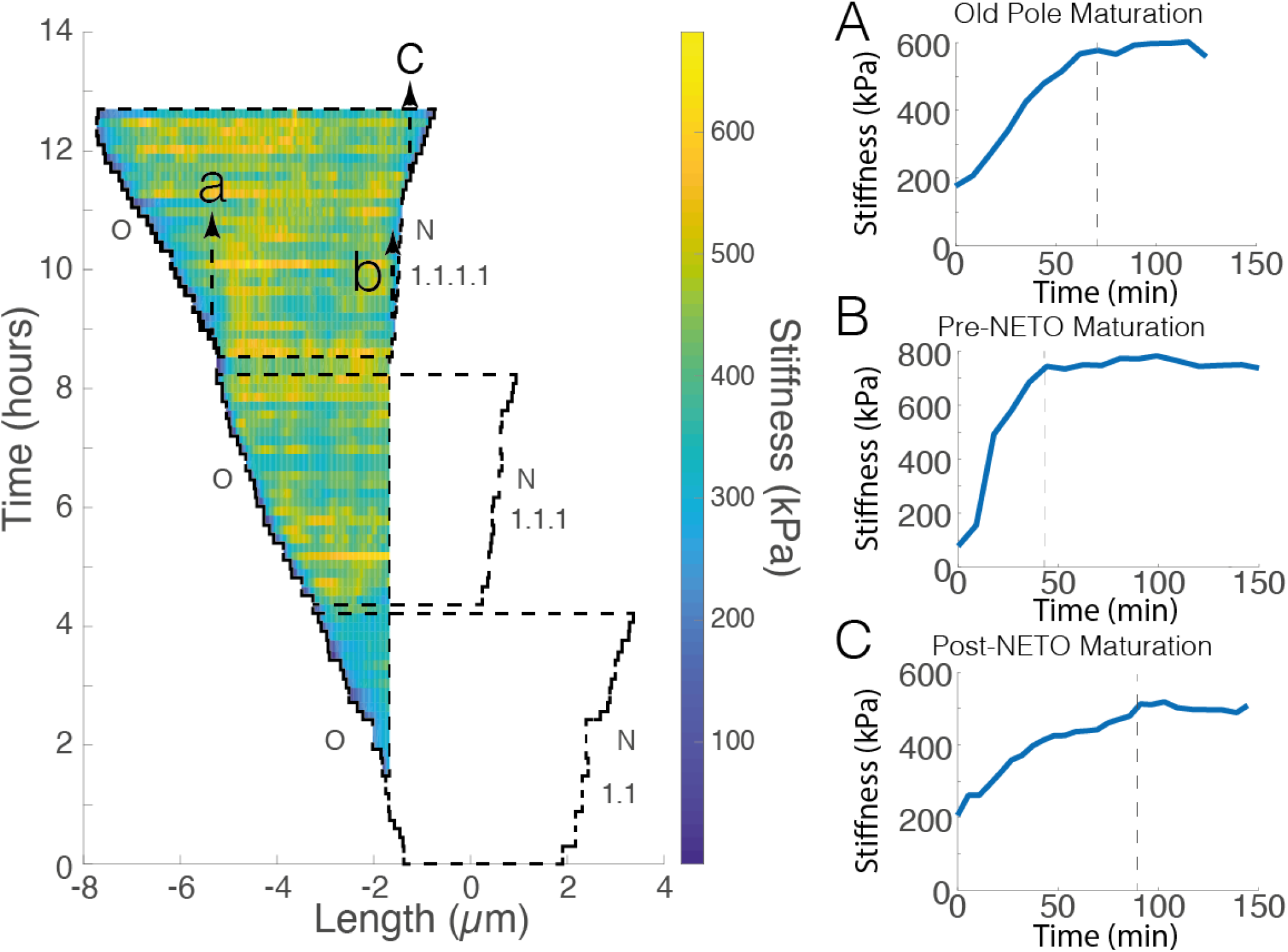
Mycobacterial cell surface mechanical growth dynamics. Kymograph representation of the cell surface of *M. smegmatis* over three successive generations of growth and division. Depicted is the spatial inheritance of cell surface material from when it is first deposited as a result of polar elongation. **(A – C)**, The dynamic of “mechanical maturation” of nascent material increases in surface stiffness until it plateaus. The rate of maturation is distinguishable depending on the age of the pole and the timing of “new-end take-off” at the new pole (*7*). The spatial mobility of mechanically mature material is static, as is equally reported for chemically distinct layers (*33*).

**Fig. S2.**
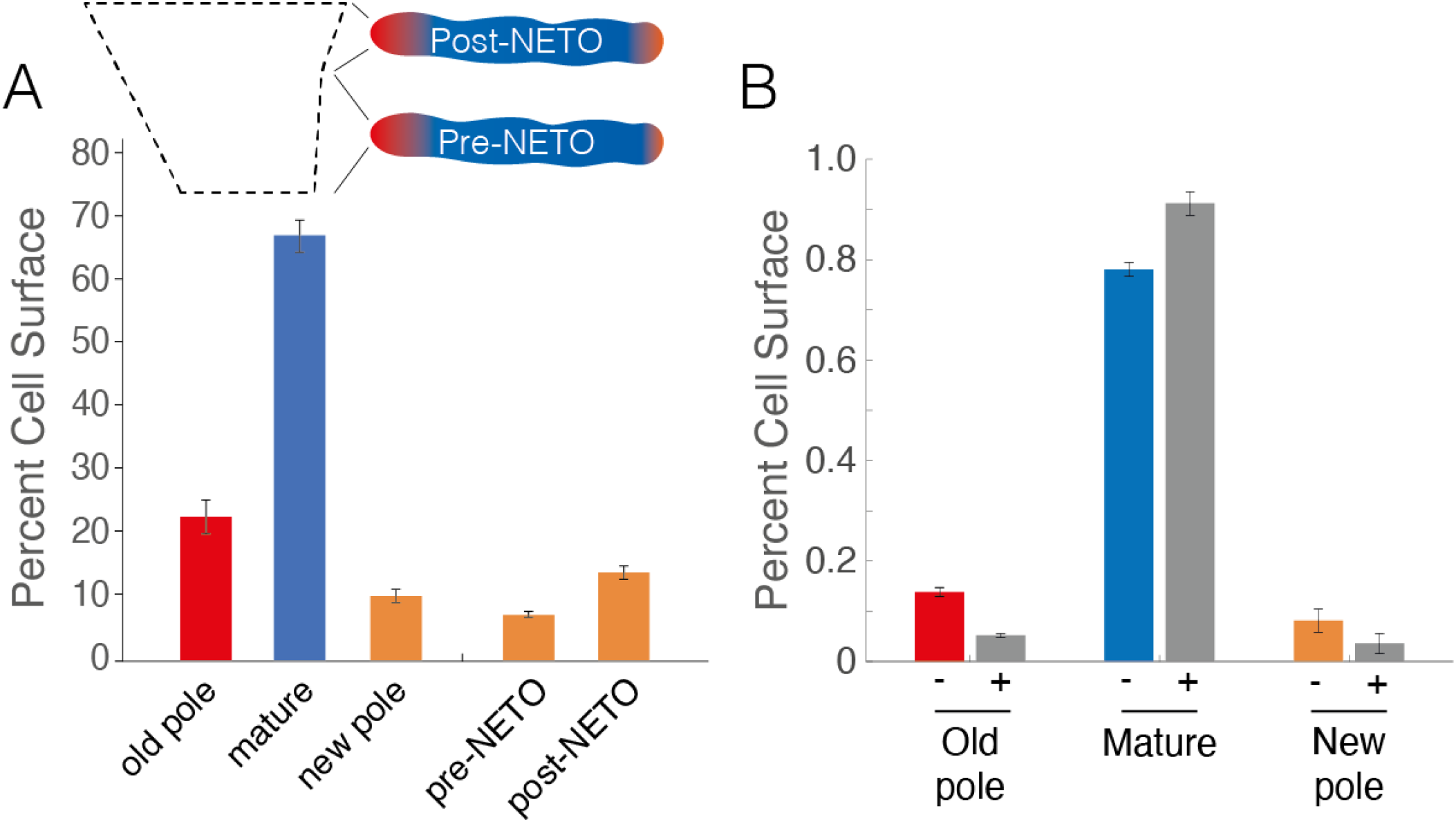
The spatial percentage of the cell surface of *M. smegmatis* as measured by LTTL-AFM quantitative nanomechanical mapping. **(A)** In growing *M. smegmatis* in axenic conditions of growth, mechanically stable cell surface material represents >60% of the cell surface (blue bar). Cell surface material that is actively undergoing an increase in mechanical rigidity represents ∼30% of the cell surface split between the old and new poles (red and orange bars). The spatial representation of new pole material can be distinguished as a function of whether new-end take-off (NETO) has taken place (*7*). **(B)** Concomitant with cessation of elongation, INH-treated bacilli (grey bars) exhibit decreased spatial distributions of mechanically nascent cell surface material at each of the cell poles. Bars represent mean +/- SEM.

**Fig. S3.**
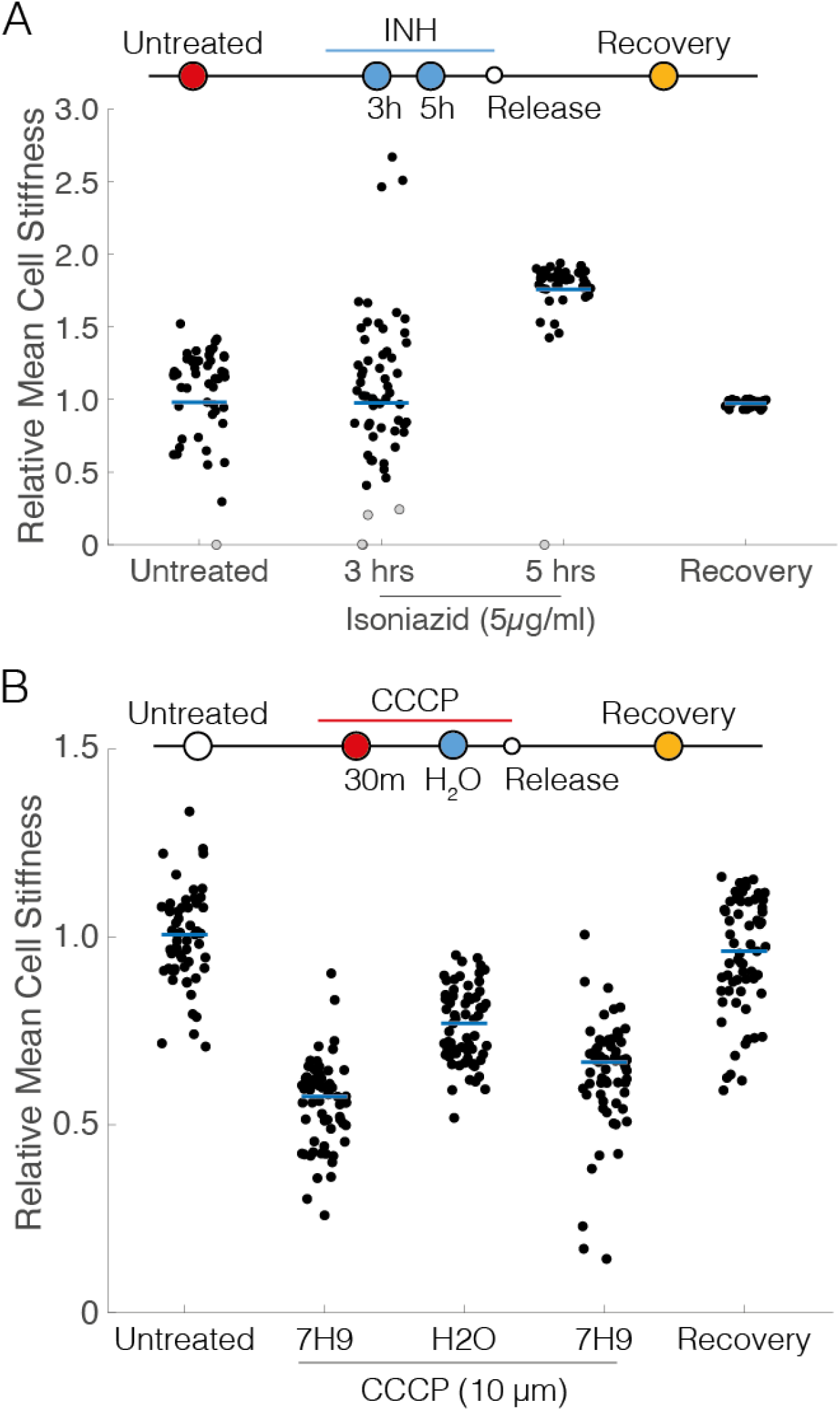
Relative mean cell surface stiffness of antibiotic-treated *M. smegmatis*. *M. smegmatis* treated with INH (5 µg/ml) **(A)** or CCCP (10 µM) **(B)**. Bacilli treated with CCCP was probed within 30 minutes; CCCP-treated bacilli were subsequently exposed to osmotic shock by transiently exchanging growth medium (7H9) with dH_2_O, before adding back 7H9 with CCCP, before later releasing bacilli from CCCP-treatment. Recovery represents the mean cell surface stiffness of cells surviving antibiotic treatment, which undergo regrowth and culminate in division, at which point the cell stiffness is probed. The relative mean cell surface stiffness represents a comparison of the mean cell surface stiffness for INH-treated cells to untreated cells. Black dots represent living cells (A and B). Grey dots represent dead cells (A).

**Fig. S4.**
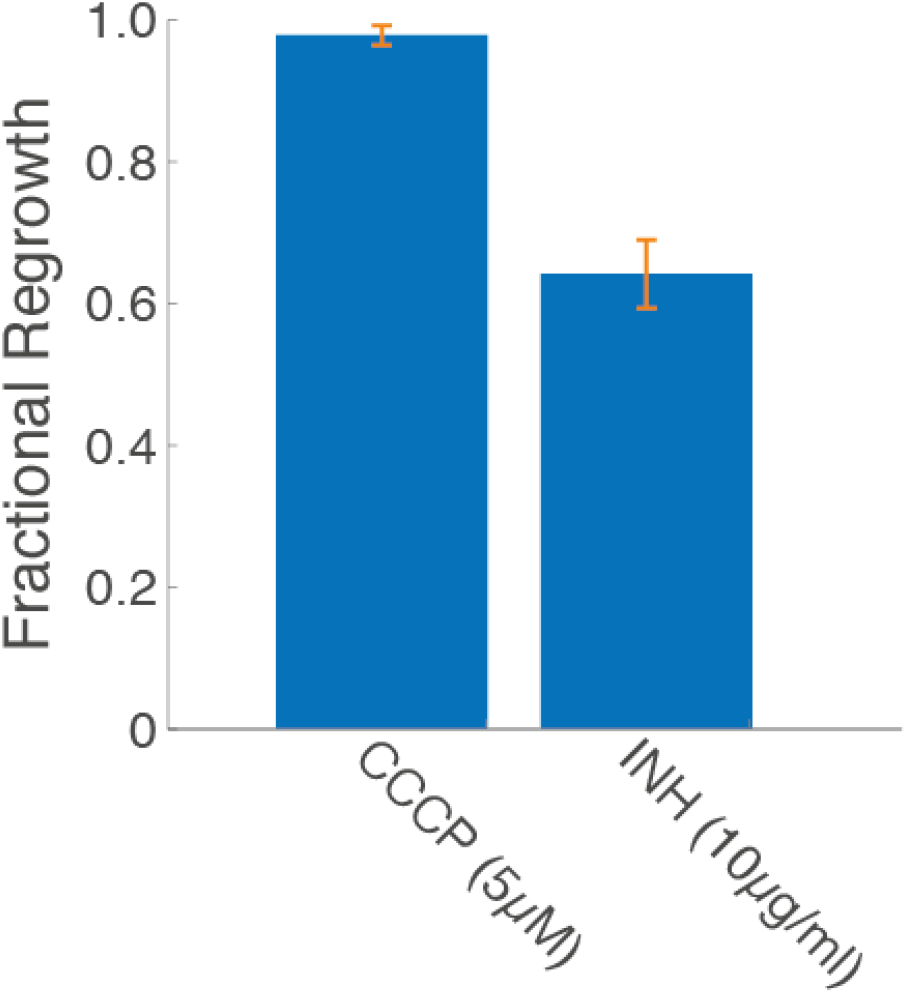
Fractional recovery of *M. smegmatis* released from antibiotic treatment. *M. smegmatis* bacilli, released from antibiotic treatment during LTTL-AFM imaging at 37°C, are counted for their ability to regrow among all surviving bacteria. Bars represent mean +/- SEM.

**Fig. S5.**
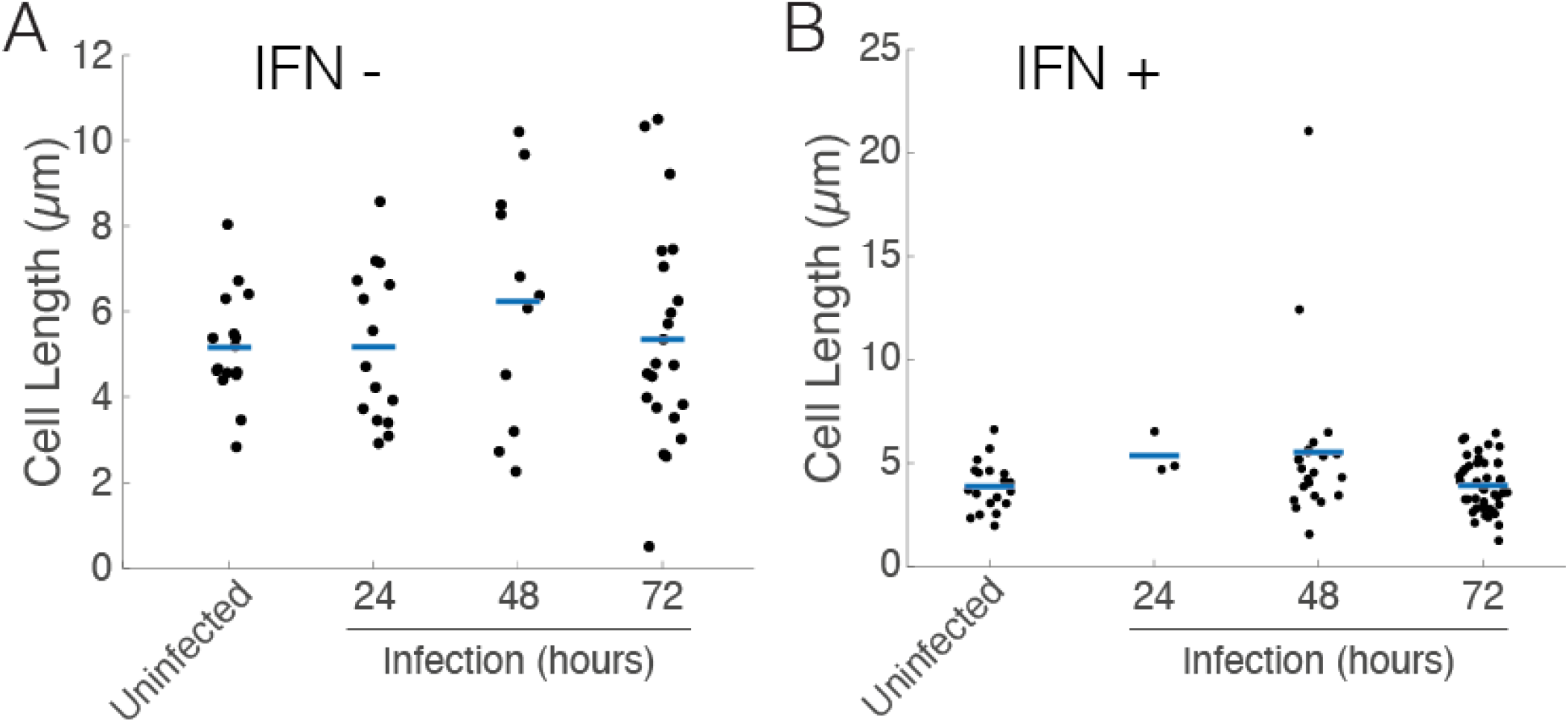
Mycobacterial cell length heterogeneity. The length of *M. smegmatis* cells is isolated and measured by AFM at room temperature (25°C) to probe the mechanical properties in static conditions. Mycobacteria were isolated from macrophages (-/+ IFNγ) and resuspended in 7H9. Blue bars represent the mean of each sample distribution.

**Fig. S6.**
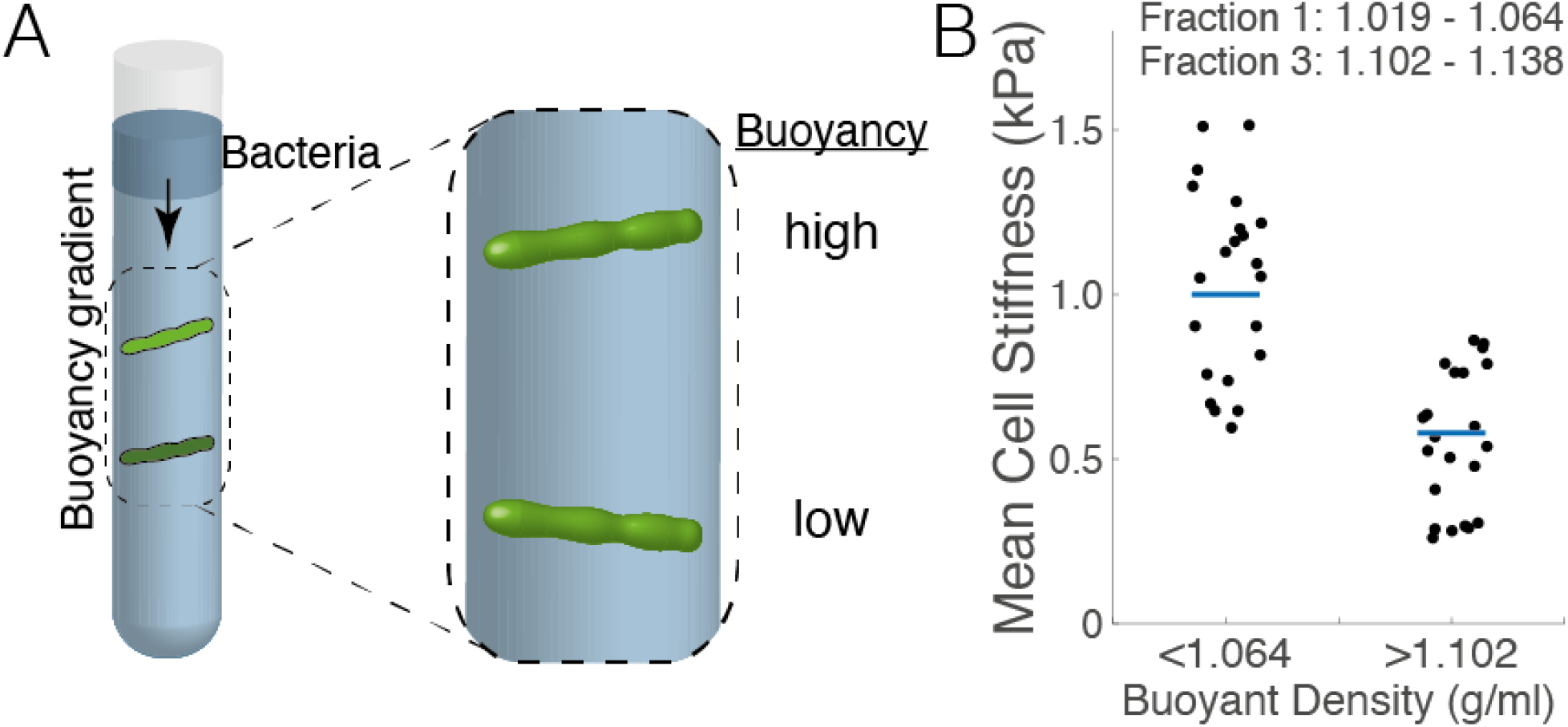
Buoyancy centrifugation as a surrogate measure for cell surface stiffness measured by AFM. **(A)** Schematic representation of buoyant density centrifugation to fractionate high and low buoyancy mycobacteria through stock isotonic Percoll (SIP). The enrichment process consists of three successive rounds of buoyancy centrifugation followed by regrowth of high and low buoyancy fractions in permissive growth medium. **(B)** AFM stiffness measurements of mycobacteria isolated from high (< 1.064 g/ml) and low (> 1.102 g/ml) buoyancy fractions. Blue bars represent mean. Black dots represent individual bacilli of *M. smegmatis*.

**Fig. S7.**
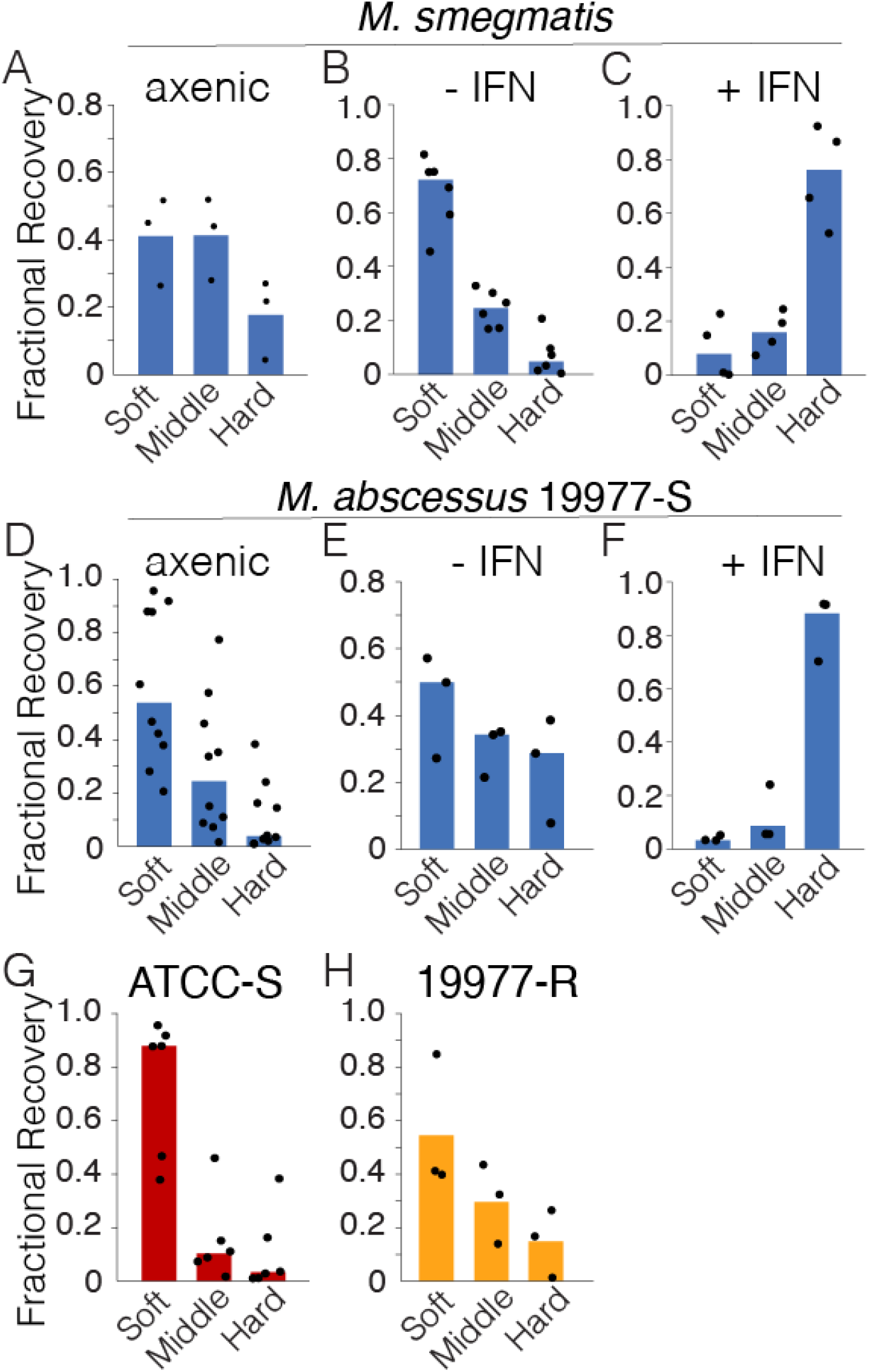
Distribution of buoyancy of mycobacteria. *M. smegmatis* **(A – C)** and *M. abscessus* strain 19977 (smooth morphotype) **(D – F)** fractionated following culturing in 7H9 growth medium (A and D) or isolated from macrophages untreated (B and E) or IFNγ-treated (C and F). Fractionation of *M. abscessus* strain ATCC (smooth morphotype) (G) and strain 19977 (rough morphotype) (H) grown in 7H9 growth medium. *M. abscessus* 19977-R represents a rough colony morphotype whereas 19977-S and ATCC-S both represent smooth colony morphotype strains. Bars represent mean and dots represent individual experimental replicates.

**Fig. S8.**
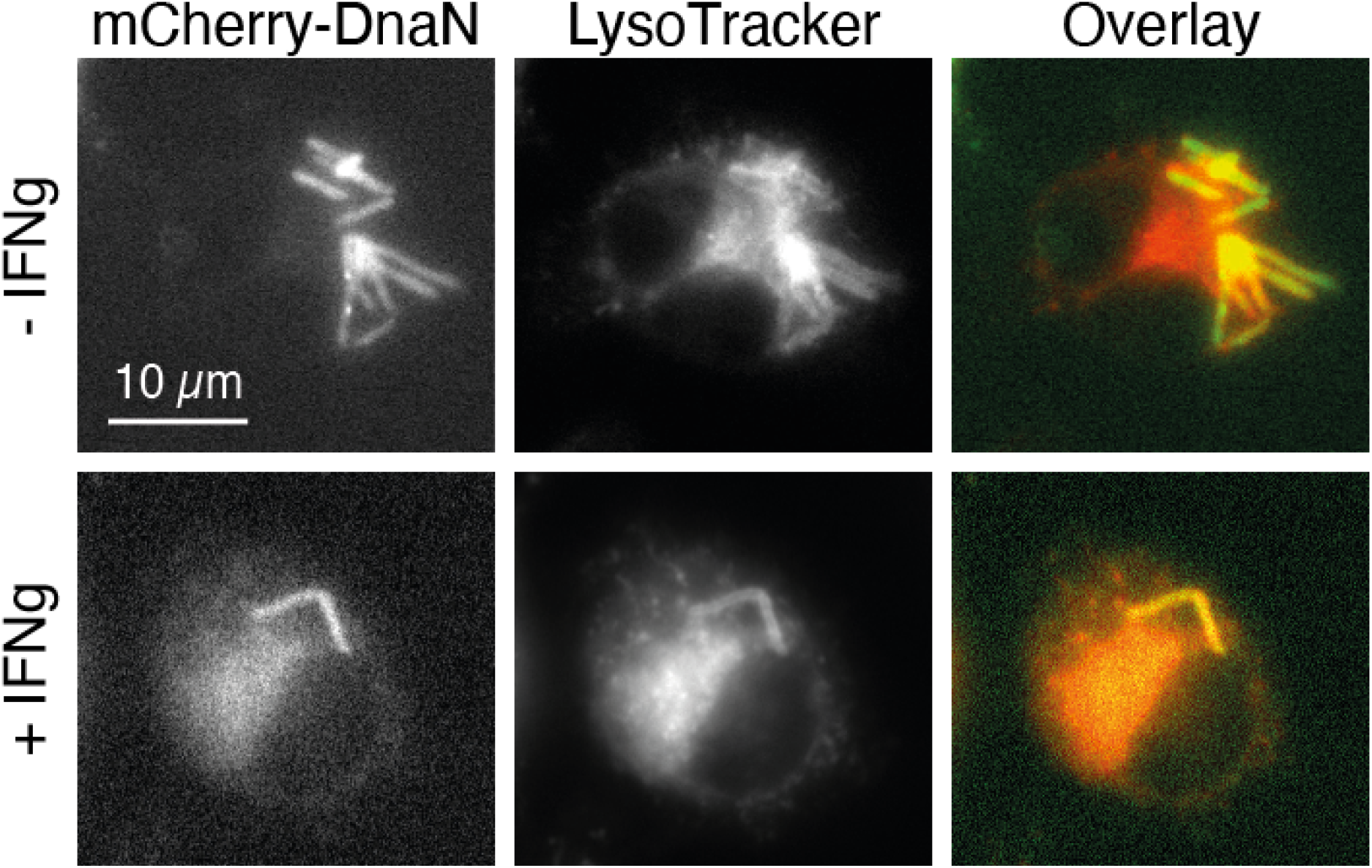
Acidification of *Mycobacterium*-containing vacuoles during infection. Optical fluorescence imaging of macrophages −/+ IFNγ harboring *M. smegmatis* expressing mCherry-tagged DnaN (integrated at the *attB* locus) (*34*). Macrophages stained with lysotracker reveal the state of acidification of vacuoles harboring *M. smegmatis* in both unstimulated and cytokine-stimulated macrophages, suggesting that vacuolar acidification is not sufficient to drive mechanical morphotype selection.

**Fig. S9.**
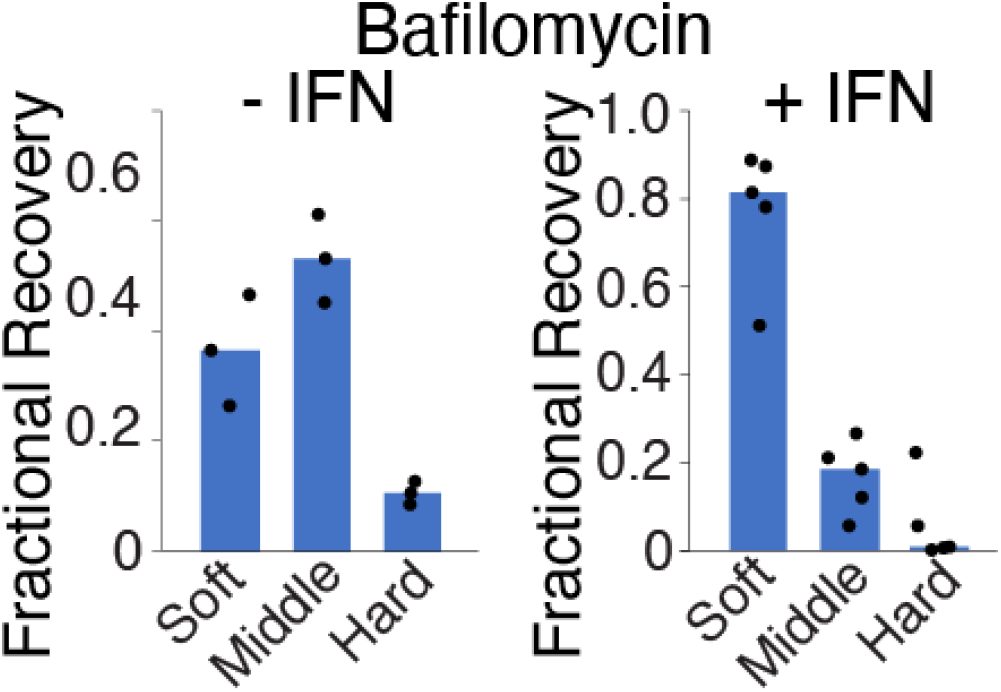
Vacuolar acidification is necessary for the selection of “high” buoyancy *M. smegmatis* in IFNγ-stimulated macrophages. Fractional recovery of *M. smegmatis* following buoyancy centrifugation after bacilli were isolated from −/+ IFNγ-treated macrophages equally treated with Bafilomycin A (10 µM, added at infection start). Bars represent mean and dots represent individual experimental replicates.

**Fig. S10.**
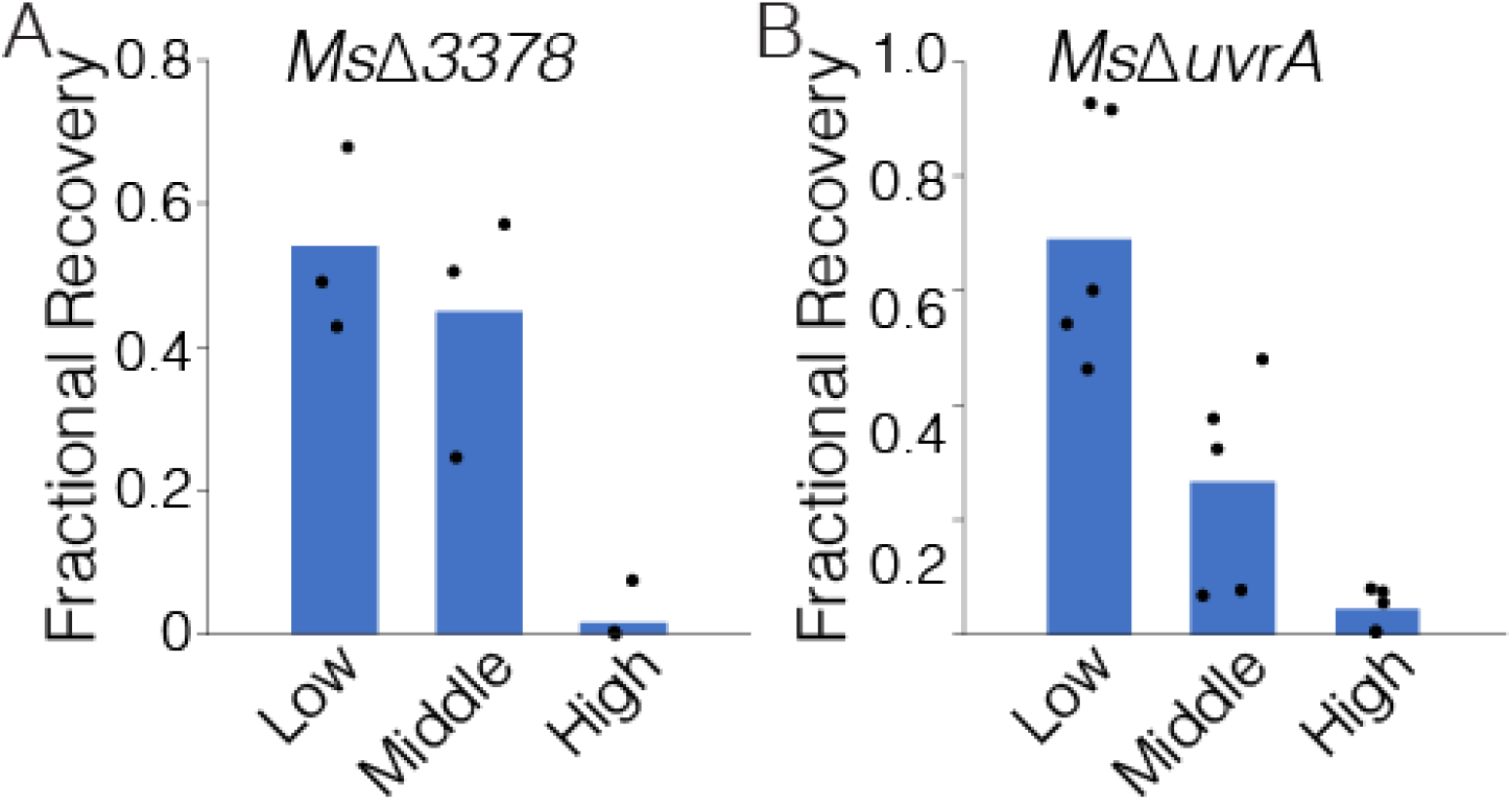
Fractional recovery of buoyancy fractionated “soft” mechanical morphotype mutants cultured in axenic conditions of growth. Bars represent mean and dots represent individual experimental replicates.

**Fig. S11.**
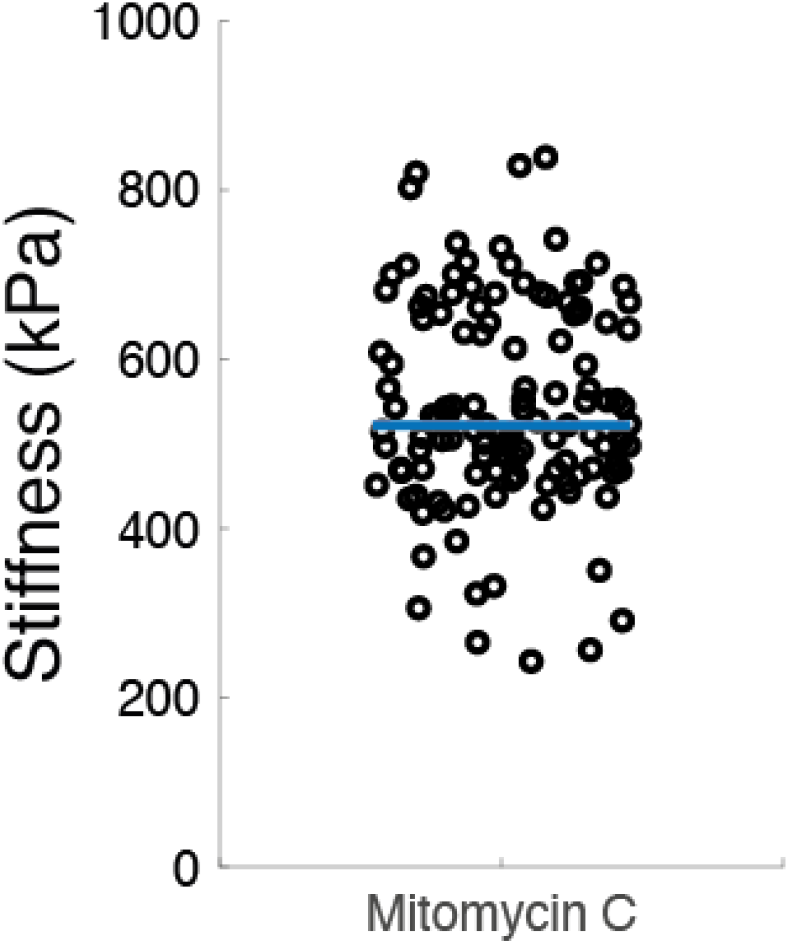
Mean cell surface stiffness of *M. smegmatis* bacilli treated with Mitomycin C. AFM imaging was used to measure the mean cell surface stiffness of *M. smegmatis* treated with Mitomycin C.

**Fig. S12.**
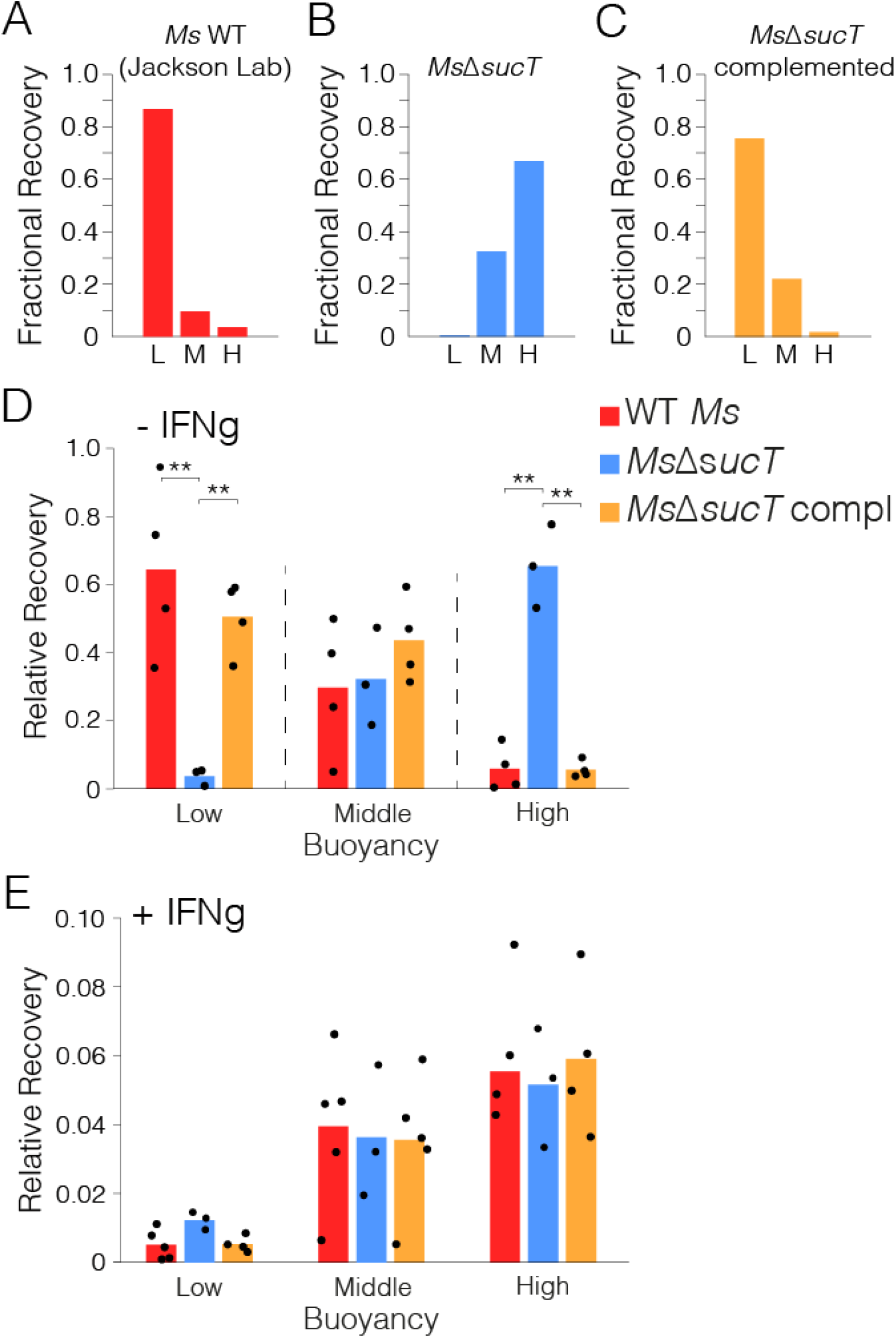
Relative fractional recovery of *M. smegmatis* Δ*sucT*. **(A – C)**, The fractional recovery of *M. smegmatis* wildtype (isolate from Colorado State University, “Jackson lab”), Δ*sucT*, and complemented *sucT* cultured in axenic conditions of growth. **(D** and **E)** Relative fractional recovery of mycobacterial strains isolated from macrophages and buoyancy fractionated. Relative fractional recovery equally shows the relative recovery of samples from macrophages. Bars represent mean and dots represent individual experimental replicates. \**P* < 0.05, \*\**P* < 0.01 by Student’s T test.

**Fig. S13.**
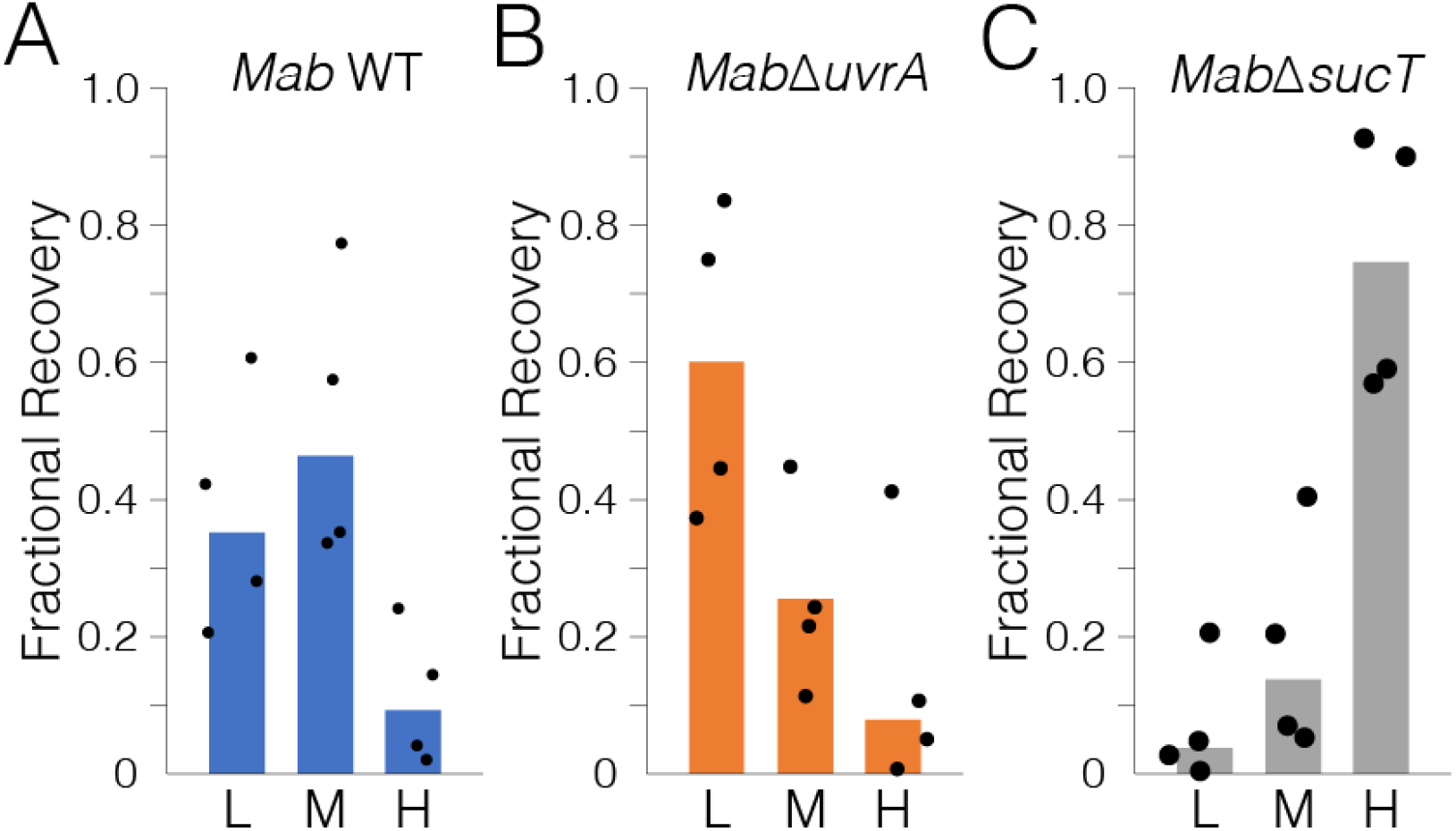
Fractional recovery of *M. abscessus* mechanical morphotype mutant candidates. buoyancy fractionated following culturing in axenic conditions of growth. Bars represent mean and dots represent individual experimental replicates.

**Fig. S14.**
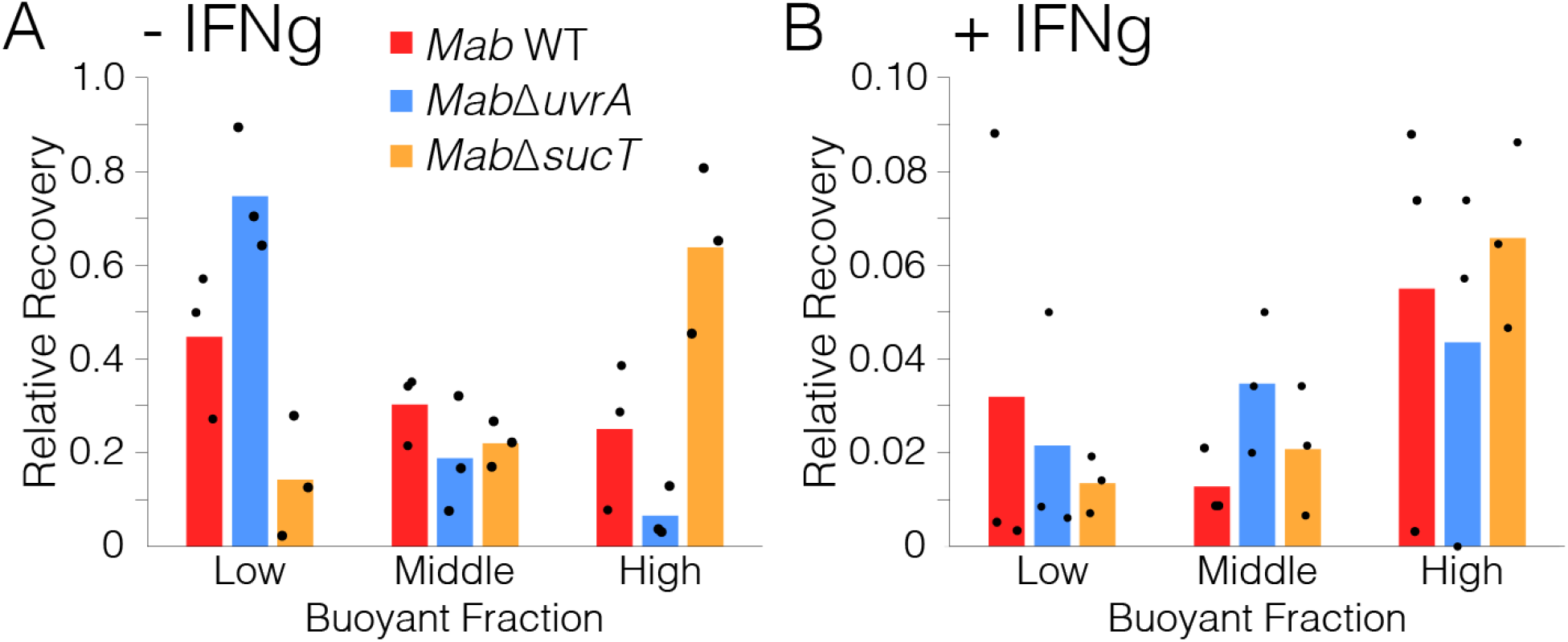
Distribution in buoyancy for “soft” and “hard” mechano-morphotype mutant candidates following infection of macrophages. Bars represent mean and dots represent individual experimental replicates.

**Fig. S15.**
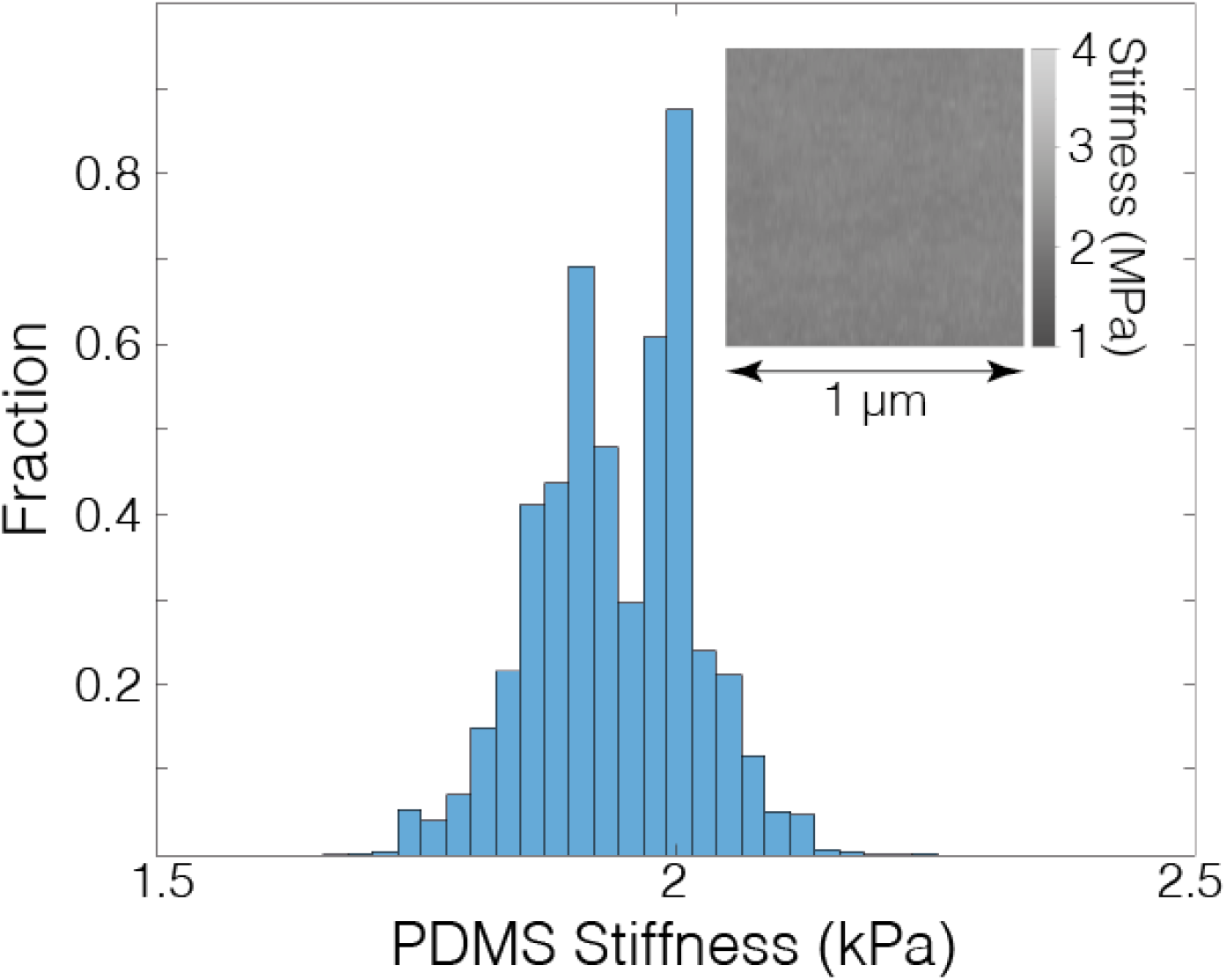
Calibrating AFM stiffness measurements from DMT modulus images was conducted using the polydimethylsiloxane-coated coverslip sample surface with known Young’s modulus of ∼1.8 MPa. Histogram depicts the distribution of measurements. Inset AFM DMT modulus image represents the measurements made using the AFM imaging mode: peak force quantitative nanomechanical mapping.

**Movie S1. LTTL-AFM imaging of *M. smegmatis* cultured in axenic conditions of growth.** AFM imaging was conducted using peak-force off-resonance tapping in time lapse. 3D-Height images are overlayed with the DMT Modulus. *M. smegmatis* was grown at 37°C and the frequency of images taken every 9 minutes. Images represent 15 µm by 7.5 µm of space. The scale of DMT modulus spans 0 – 3 MPa. See Figure 1b and Supplementary Figure 16 for representative schematic images of the same time-lapse.

**Movie S2. LTTL-AFM imaging of *M. smegmatis*** Δ***ldtAEBCG*+*F* cultured in axenic conditions of growth.** AFM DMT modulus images depicting the cell surface stiffness of a mycobacterial mutant in which all L, D Transpeptidases are absent resulting in less peptidoglycan crosslinking at the new pole defective. The mechanical consequence is bulging spatially localized near the new pole, which happens at a rate that is controlled by the addition of new material at the cell wall (*1*).

**Movie S3. LTTL-AFM imaging of *M. smegmatis* WT cells cultured in axenic conditions of growth and antibiotic stress.** AFM peak force error images depict the cell surface of *M. smegmatis* cells.

**Movie S4. LTTL-AFM imaging of *M. smegmatis* WT cells cultured in axenic conditions of growth and antibiotic stress as per Supplementary Video 3.** AFM DMT Modulus images depict the cell surface of *M. smegmatis* cells in conditions of growth, treatment with the bacteriostatic CCCP (5 µM), and recovery of bacilli following washout of the drug.

